# AR-V7 condensates drive androgen-independent transcription in castration resistant prostate cancer

**DOI:** 10.1101/2025.01.08.631986

**Authors:** Shabnam Massah, Nicholas Pinette, Jane Foo, Sumanjit Datta, Maria Guo, Robert Bell, Anne Haegert, Tahsin Emirhan Tekoglu, Mailyn Terrado, Stanislav Volik, Stephane Le Bihan, Jennifer M. Bui, Nathan A. Lack, Martin E. Gleave, Suhn K. Rhie, Colin C. Collins, Jörg Gsponer, Nada Lallous

**Affiliations:** Vancouver Prostate Centre, University of British Columbia, Vancouver, BC, Canada; Department of Medical Pharmacology, School of Medicine, Koç University, Istanbul, 34450, Turkey; Michael Smith Laboratories, University of British Columbia, Vancouver, BC, Canada; Department of Biochemistry and Molecular Medicine and the Norris Comprehensive Cancer Center, Keck School of Medicine, University of Southern California, Los Angeles, CA 90089, USA

## Abstract

Biomolecular condensates organize cellular environments and regulate key processes such as transcription. We previously showed that full-length androgen receptor (AR-FL), a major oncogenic driver in prostate cancer (PCa), forms nuclear condensates upon androgen stimulation in androgen-sensitive PCa cells. Disrupting these condensates impairs AR-FL transcriptional activity, highlighting their functional importance. However, resistance to androgen deprivation therapy often leads to castration-resistant prostate cancer (CRPC), driven by constitutively active splice variants like AR variant 7 (AR-V7). The mechanisms underlying AR-V7’s role in CRPC remain unclear. In this study, we characterized the condensate-forming ability of AR-V7 and compared its phase behavior with AR-FL across a spectrum of PCa models and *in vitro* conditions. Our findings indicate that cellular context can influence AR-V7’s condensate-forming capacity. Unlike AR-FL, AR-V7 spontaneously forms condensates in the absence of androgen stimulation and functions independently of AR-FL in CRPC models. However, AR-V7 requires a higher concentration to form condensates, both in cellular contexts and *in vitro*. We further reveal that AR-V7 drives transcription via both condensate-dependent and condensate-independent mechanisms. Using an AR-V7 mutant incapable of forming condensates, while retaining nuclear localization and DNA-binding ability, we reveal that the condensate-dependent regime activates part of the oncogenic KRAS pathway in CRPC models. Genes under this condensate-dependent regime were found to harbor significantly higher numbers of AR-binding sites and exhibited boosted expression in response to AR-V7. These findings uncover a previously unrecognized role of AR-V7 condensate formation in driving oncogenic transcriptional programs and shed light on its unique contribution to CRPC progression.

**Highlights:**

- AR-V7 condensates form independently of both androgens and AR-FL in CRPC models.
- AR-V7 mediates condensate-dependent and independent transcription
- Condensate-dependent transcription enables boosted expression of oncogenic KRAS genes
- Condensate-dependent genes exhibit an exponential increase in expression, with a higher number of AR binding sites potentially playing a key role in their reliance on condensate formation.

*Graphical Abstract:* 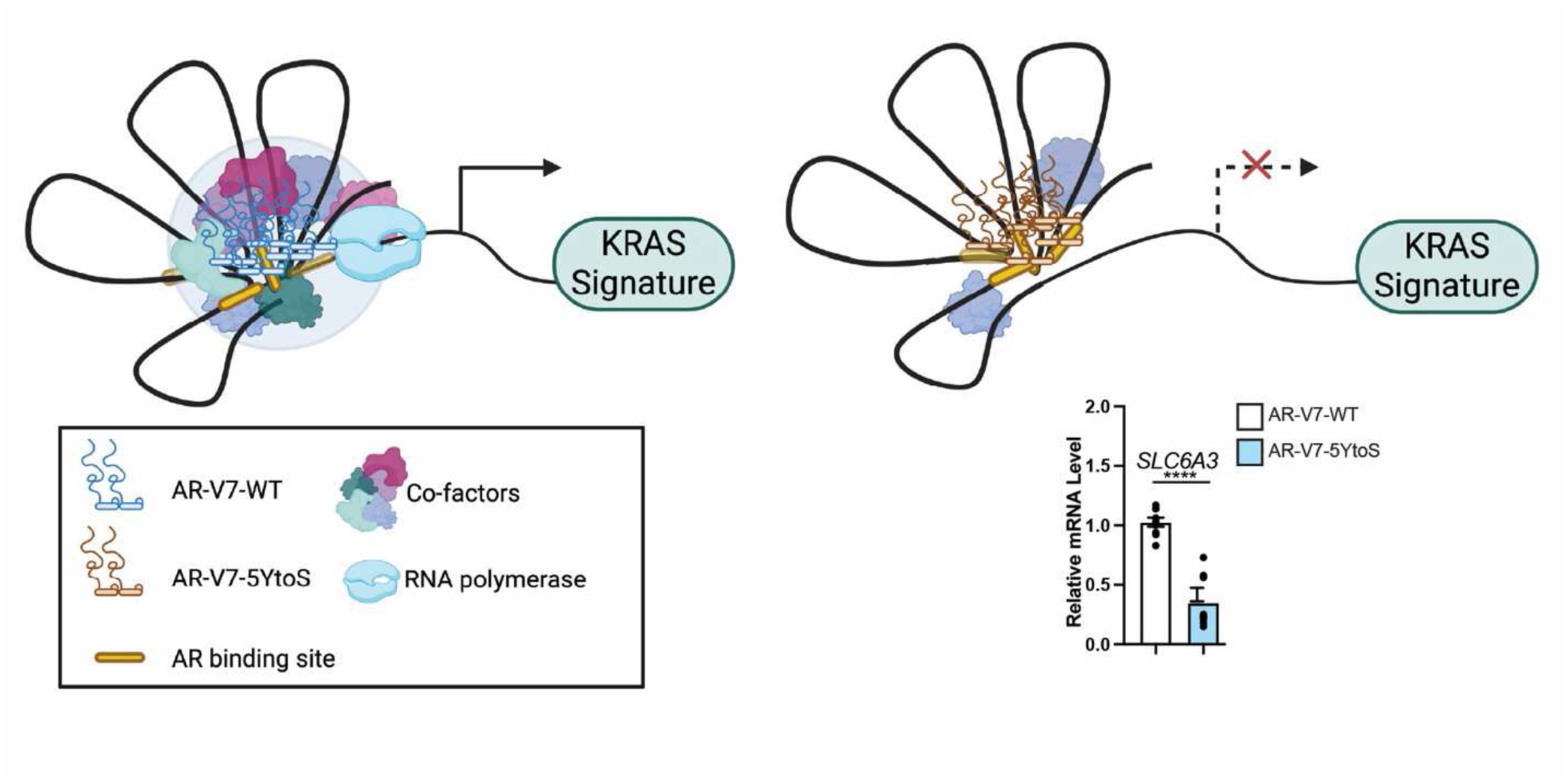

## Introduction

Biomolecular condensates are membraneless structures that compartmentalize the intracellular environment, creating confined spaces for specific cellular processes, including epigenetic regulation, stress response, transcription, and DNA repair ^1–5^. These condensates form through phase separation, enabling dynamic and reversible formation of the confined spaces ^6^. Many transcription-related proteins, such as transcription factors (KLF4 ^7^, FOXA-1 ^8^, OCT4 ^3^, ER ^3^, GR ^9^, MYC ^10^), transcriptional coactivators (MED1 ^4^, BRD4), RNA polymerase II (RNA-Pol II^4^), and epigenetic regulators (p300 ^11^, HP1 ^12^), are known to form condensates that facilitate the transcriptional process. The ability of these proteins to form condensates often depends on intrinsically disordered regions (IDRs) within their sequences ^13–16^ . Moreover, these IDRs play a key role in selectively recruiting specific proteins into the condensates while excluding others ^17^. For instance, the IDR of MED1 (MED1-IDR) selectively partitions with the IDRs of co-regulators like CTR9 and SPT6, which share similar alternating patterns of charged amino acids, while excluding proteins such as FUS and PTBP1, whose IDRs lack such charge patterning ^17^. This specificity in partitioning underscores the selective interactions that govern condensate composition, likely playing a crucial role in shaping the functional outcomes of transcription.

We recently demonstrated that the full-length androgen receptor (AR-FL), a ligand-activated transcription factor, phase separates *in vitro* and forms dynamic biomolecular condensates upon androgen stimulation in androgen-sensitive prostate cancer (PCa) models ^18^. AR-FL condensates form at enhancers of AR target genes, such as *FKBP5*, and colocalize with key transcriptional machinery, including RNA polymerase II (RNA Pol II) and the coactivator MED1^18^. Importantly, AR transcriptional activity in PCa models follows changes in cellular condensate content induced chemically or by cofactor silencing ^18^. These and findings from other labs ^13, 19–22^ have highlighted the functional importance of AR condensates in orchestrating androgen-induced transcriptional activity.

Dysregulation of biomolecular condensates has increasingly been linked to a wide range of diseases, including cancer. In the case of the oncoprotein MYC, for instance, high expression levels in tumors promote excessive condensate formation and drive aberrant transcriptional activity^10^. As AR is the main oncogenic driver in PCa, it has been speculated that enhanced compartmentalization of AR and coactivators via phase separation may also play an important role in the activation of oncogenic transcription programs in PCa. Of particular interest in this context is the AR variant V7 (AR-V7) ^23, 24^ that lacks the C-terminal ligand-binding domain (LBD), where androgens bind to activate the AR, making this variant constitutively active. AR-V7 expression is rarely observed in primary PCa ^25^. However, its expression increases in castration resistant PCa (CRPC) ^26^, a lethal form of the disease that becomes androgen-independent ^27–31^ following treatment with AR pathway inhibitors such as abiraterone acetate or enzalutamide ^32^. Intriguingly, the majority of CRPC patients remain reliant on AR activity, facilitated by various mechanisms including the expression AR-V7. How AR-V7 contributes to this aggressive form of PCa and whether condensate formation plays a role, remains largely elusive.

Previous studies have shown that AR-V7 forms condensate-like foci in COS7 or PC3 cells ^20, 33^. However, we found that removing the structured LBD inhibits the ability of the AR to form condensates in LNCaP cells^18^. Moreover, we found that the N-terminal domain (NTD) of the AR, an IDR that plays a key role in recruiting coactivators essential for transcriptional regulation^34^, is unable to form condensates in LNCaP cells on its own^18^. Taken together, these findings suggest that the cellular context may influence the ability of AR-V7 to form condensates, highlighting the need for a deeper understanding of the mechanisms underlying their formation in cancer cell environments. Here, we investigated AR-V7 condensate formation across PCa models and its role in activating transcriptional programs associated with the CRPC phenotype. We reveal that AR-V7 forms condensates independently of androgen stimulation and AR-FL in CRPC models, but requires chromatin accessibility and DNA binding for this to happen. Importantly, we identified both condensate-dependent and -independent transcriptional paradigms regulated by AR-V7. Condensate-dependent genes showed significant enrichment for KRAS pathway genes ^35–37^. Furthermore, condensate dependent genes exhibited exponential expression and harbored an increased number of binding sites.

## Results

### AR-V7 forms foci independently of androgens

We previously reported that AR-FL forms nuclear foci with liquid-like properties when stimulated by androgens in androgen-sensitive LNCaP cells, whereas AR-V7 does not ^18^. Therefore, we first evaluated whether cellular context affects the ability of endogenous AR-V7 to form such foci. Specifically, we used immunofluorescence microscopy to monitor foci formation, both in the presence and absence of dihydrotestosterone (1 nM DHT), in CRPC models that endogenously express AR-V7 (LN95, 22Rv1 and VCaP) using an AR-V7 specific antibody (RevMab 31-1109-00). We found that endogenous AR-V7 forms foci independently of androgens in these PCa models (Figure 1A-B and Figure S1A-B). Additionally, we found that endogenous AR-FL, detected by a C-terminal antibody (Santa Cruz sc-815), translocates to the nucleus and forms foci only after stimulation with androgens (Figure 1C-D), confirming our previously reported behaviour for AR-FL in other PCa models ^18^. To further evaluate the impact of the cellular environment on foci formation, we assessed the ability of exogenous AR-V7 and AR-FL, tagged with a non-dimerizing Enhanced Green Fluorescent Protein (mEGFP), to form foci in various PCa models including AR positive models (LNCaP, LN95, 22Rv1, VCaP) and an AR negative one (PC3). We transfected these PCa models with either AR-FL-mEGFP or AR-V7-mEGFP and quantified foci in the absence and presence of androgens. To account for variation in transfection efficiency, we normalized foci density by the nuclear expression levels of mEGFP-tagged proteins. We found that exogenous AR-V7-mEGFP forms foci in CRPC models LN95, 22Rv1, and VCaP independently of androgen stimulation, but rarely in the androgen-sensitive LNCaP cells (Figure 1E-F). As LN95 cells are closely related to LNCaP and were generated through long-term culture in androgen-deprivation conditions^38^, we assessed whether the change in AR-V7-mEGFP behavior (Figure 1E-F) is due to differences in expression levels between the two models. However, we found similar expression levels of AR-V7 between LNCaP and LN95 cells (Figure S1C-D) and observed that the average nuclear intensity of AR-V7-mEGFP was comparable between them (Figure S1E), suggesting that cellular context potentially influences foci density. Regarding AR-FL, we found that, similar to the endogenous receptor, exogenous AR-FL-mEGFP required androgen stimulation to translocate to the nucleus and form foci (Figure 1G-H). Interestingly, we found that both AR-FL-mEGFP and AR-V7-mEGFP form foci in the AR-negative CRPC model PC3, corroborating a previous report regarding the latter^20^ and further underscoring the influence of the cellular environment on AR-V79s ability to form foci.

**Figure 1:**
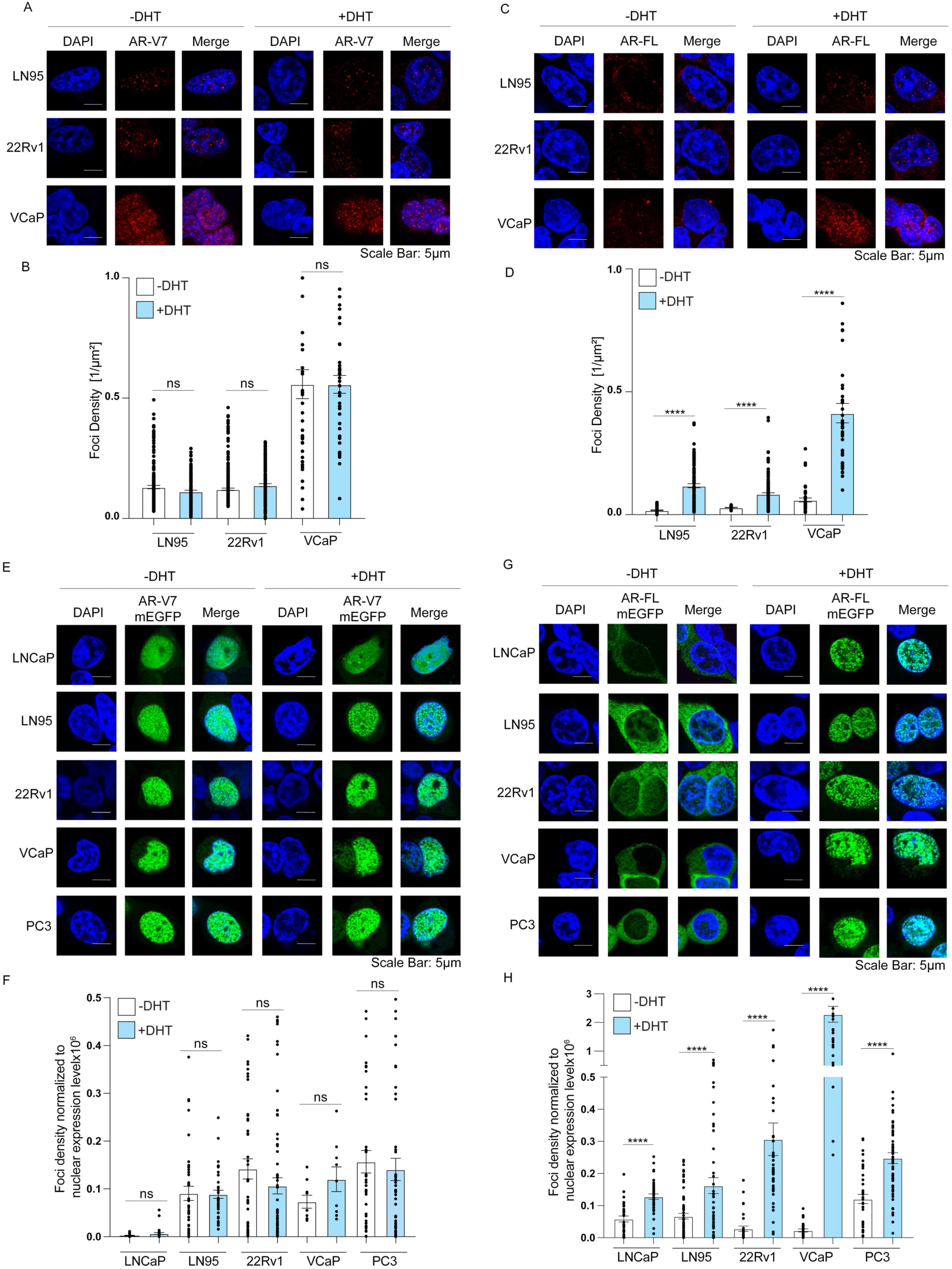
PCa models (LN95, 22Rv1, and VCaP) were serum starved for 48 hrs then stimulated with either 1 nM DHT or vehicle for 2 hrs. Endogenous AR-V7 (A-B) and AR-FL (C-D) condensates were detected by immunofluorescence using isoform specific antibody (RevMab 31-1109-00 for AR-V7 and Santa Cruz sc-815 for AR-FL). Foci densities were quantified from a total of 45 cells from three independent experiments (n= three biological replicates and 15 images per replicate) Data are presented as mean ± SEM. Statistical significance was determined using a t-test. PCa models (LNCaP, LN95, 22Rv1, VCAP, and PC3 cells) were transfected with 2 μg of either AR-FL-mEGFP or AR-V7-mEGFP in serum starved media for 48 hrs. The cells were then stimulated with either 1 nM DHT or vehicle for 2 hrs. The condensates formed by transiently expressed AR-V7-mEGFP (E-F) or AR-FL-mEGFP (G-H) were detected by confocal microscopy. Foci density were quantified from a total of 45 cells from three independent experiments and were normalized to the nuclear expression level (n= three biological replicates and 15 images per replicate). Data are presented as mean ± SEM. Statistical significance was determined using a t-test.

### AR-V7 foci form independently of AR-FL in CRPC cell models

It has been reported that AR-V7 can heterodimerize with AR-FL to transcribe target genes ^21, 39–41^. However, recent studies also show that AR-V7 can independently regulate gene expression and has a distinct transcriptome ^42–44^. We thus asked whether AR-FL expression affects AR-V79s ability to form foci in CRPC models. Our observations show that AR-FL translocation to the nucleus after androgen stimulation does not change the density of AR-V7 foci, suggesting it does not affect AR-V7 foci formation (Figure 1E-F). To further explore the interdependency between the two isoforms, we targeted AR-FL for proteasomal degradation by using the PROTAC molecule ARV110 ^45^ (Figure 2A). We thus treated 22Rv1 and LN95 cells with 100 nM ARV110 for 16 hrs in the absence and presence of androgens. As expected, we observed a significant reduction in AR-FL protein levels upon ARV110 treatment, while seeing minimal effects on AR-V7 levels, as assessed by western blot (Figure 2B and S2A). We then examined the effect of this treatment on foci formed by both endogenous AR-FL and AR-V7 in 22Rv1 and LN95 cells. Upon androgen stimulation, AR-FL forms nuclear foci, which are significantly reduced following ARV110 treatment (Figure 2C-D and S2B, C). However, the number of AR-V7 foci remains unaffected by the loss of AR-FL post ARV110 treatment (Figure 2E-F and Figure S2D-E). To further validate these findings, we co-transfected the AR-negative PC3 cells with an increasing amount of AR-V7-mEGFP plasmid (500 ng, 750 ng, and 1000 ng) in the presence of a fixed amount of unlabelled AR-FL plasmid (750 ng), then assessed the number of AR-V7 foci in the absence and presence of androgens. We found that in the absence of androgens, AR-V7 does not facilitate the nuclear translocation or foci formation of AR-FL (Figure 2G-I). We also assessed whether androgen activation of AR-FL lowers the concentration needed for AR-V7 to form foci. Our results indicate that AR-FL activation does not increase AR-V7 foci density at any of the studied concentrations or reduce the concentration needed for AR-V7 foci formation (Figure 2J-K and 2I). These findings demonstrate that AR-V7 forms foci independently of AR-FL in different CRPC models and does not promote AR-FL nuclear translocation.

**Figure 2.**
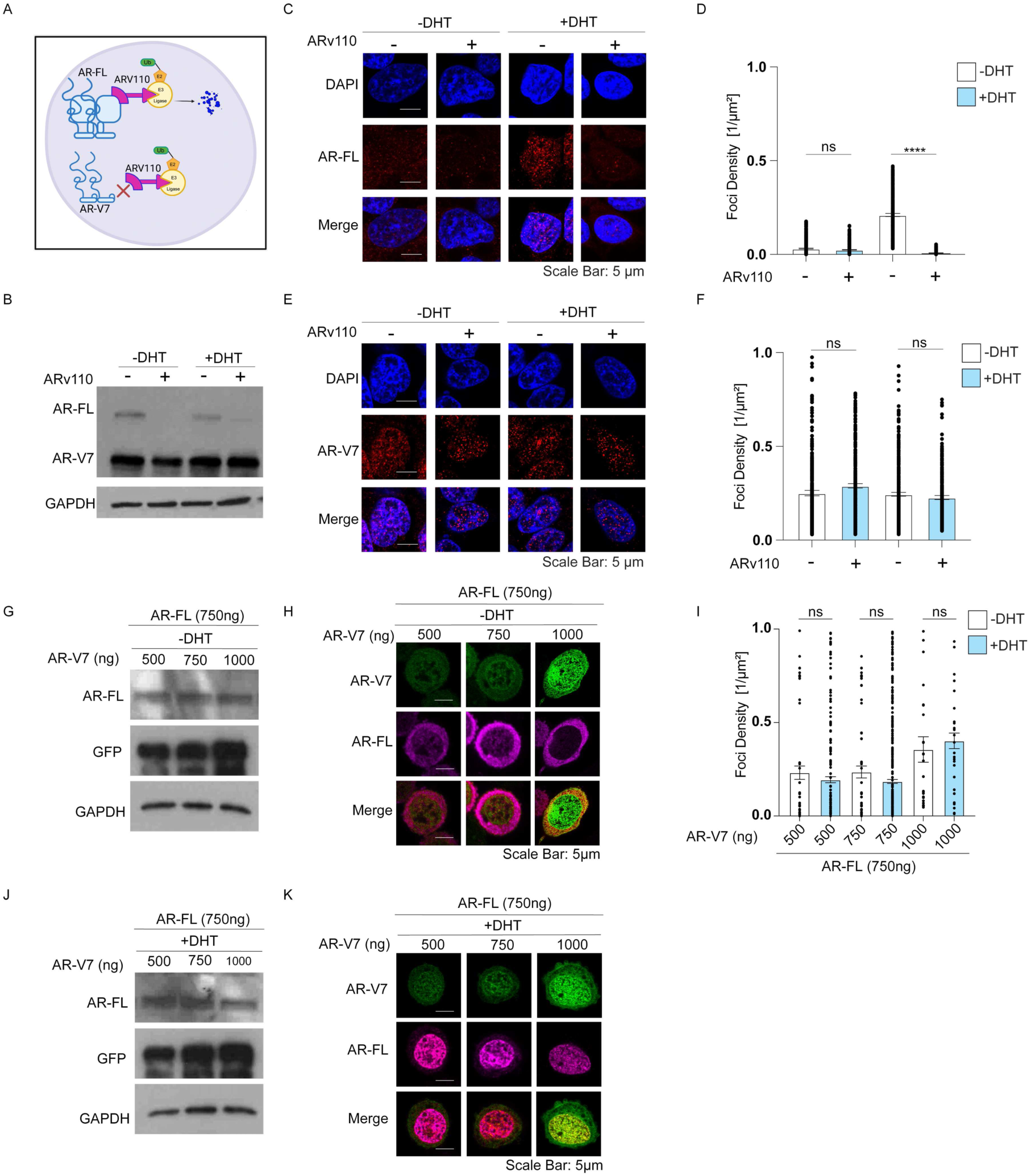
(A) Schematic illustrating the specific degradation of AR-FL by PROTAC. (B-F) 22Rv1 cells were serum-starved for 48 hrs, then treated with vehicle or 100 nM ARV110 for 16 hrs. Cells were subsequently stimulated with 1 nM DHT or vehicle for 2 hrs. (B) The degradation of AR-FL was validated by western blot using AR-441 antibody (targeting the NTD). Endogenous AR-FL (C-D) and AR-V7 (E-F) condensates were detected using isoform-specific antibodies AR-C19 (sc-815) and AR-V7 (Rev-Mab 31-1109-00), respectively. Cells were imaged by confocal microscopy and nuclear foci densities were determined (n = 3 biological replicates, 15 images per replicate). Data are presented as mean ± SEM. Statistical significance was determined using a t-test. (G-K) PC3 cells were transfected with varying concentrations of AR-V7-mEGFP in the presence of AR-FL-pCDNA (750 ng). After 48 hrs in serum-starved media, cells were stimulated with either vehicle (Figures G-H) or 1 nM DHT (Figures J-K) for 2 hours. Whole-cell lysates were analyzed for AR-V7-mEGFP levels using an anti-GFP antibody and for AR-FL levels using an isoform-specific antibody (AR C-19) (Figures G-J). AR-FL condensate formation was assessed by immunofluorescence with isoform-specific antibodies (AR C-19), and colocalization with AR-V7-mEGFP condensates was examined by confocal microscopy (Figures H-K). Quantification of foci density is presented in (I). Data are presented as mean ± SEM. Statistical significance was determined using a t-test. The original colors for AR-FL was red. For compatibility with colorblind readers, the colors were adjusted in Photoshop using the Hue setting (red shifted by -65).

### AR-V7 requires higher concentrations than AR-FL to phase separate in PCa cells and in vitro

We next investigated whether AR-V7 foci are dynamic and exhibit characteristics of liquid-like condensates. To achieve this, we transfected 22Rv1 and LN95 cells with AR-V7-mEGFP plasmid and performed fluorescence recovery after photobleaching (FRAP). We observed that AR-V7-mEGFP condensates recovered fluorescence following photobleaching on a timescale of seconds, with an average half-time of t_1/2_ = 6.4 s in 22Rv1 cells (95% CI: 5.6-7.4 s) and a mobile fraction of 43% (95% CI: 32%-54%) (Figure 3A). Similar results were obtained in LN95 cells with an average half time of 5.7 s (95% CI: 4.3-7.7 s) and mobile fraction of 48% (95% CI: 31%-56%) (Figure 3B). We further assessed the dynamic properties of AR-FL-mEGFP in 22Rv1 and LN95. In 22Rv1 cells, AR-FL-mEGFP condensates recovered after photobleaching with an average half-time of t¡/¢ **=** 7.5 s (95% CI: 6.2-9.2 seconds) and a mobile fraction of 35% (95% CI: 34%358%) (Figure 3C). Similarly, in LN95 cells, AR-FL-mEGFP condensates recovered with an average half-time of t¡/¢ = 5.9 s (95% CI: 5.0-7.1 seconds) and a mobile fraction of 40% (95% CI: 51%363%) (Figure 3D). These findings suggest that AR-V7 and AR-FL exhibit comparable dynamic properties in CRPC models.

**Figure 3.**
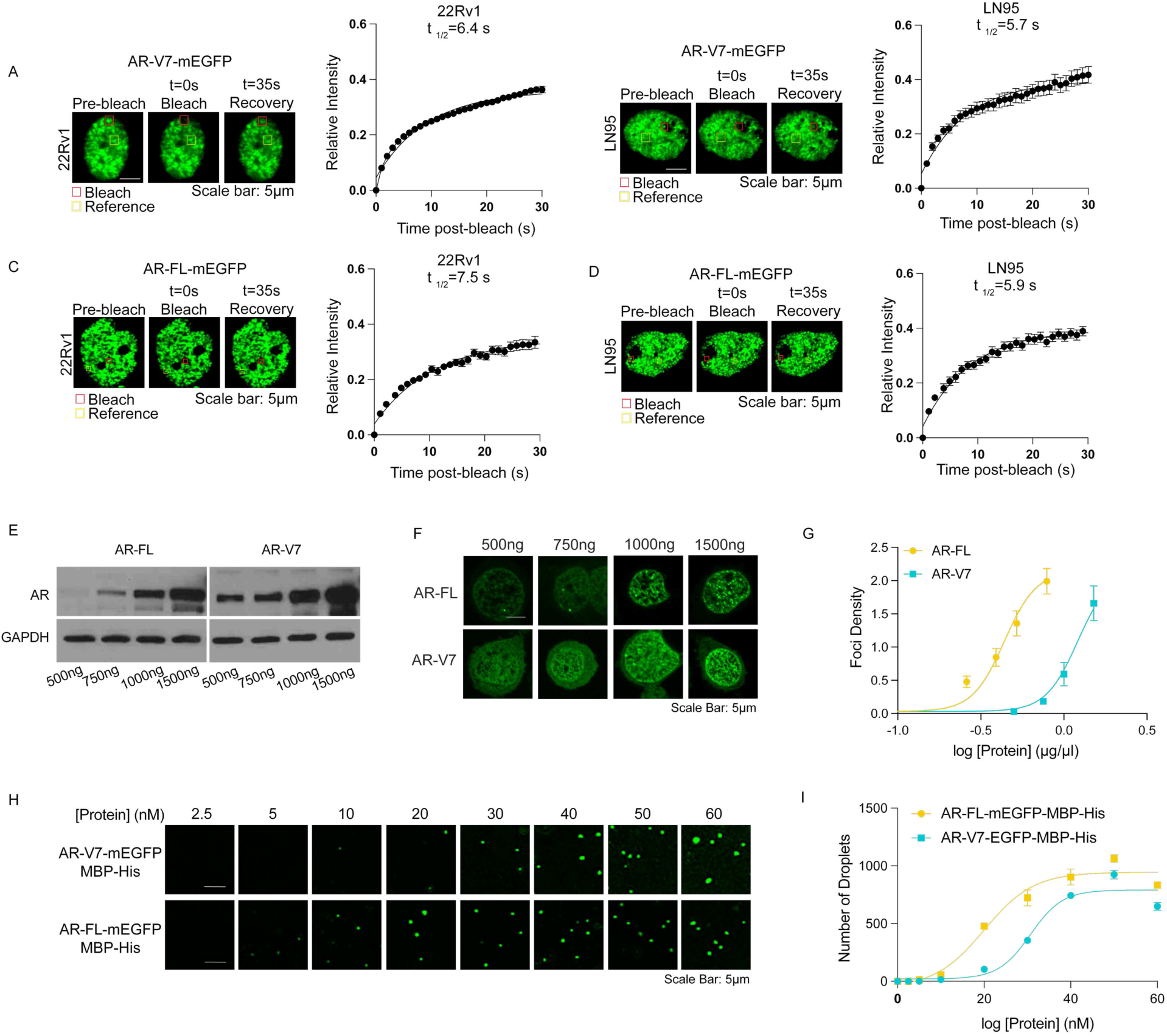
FRAP assay in 22Rv1 and LN95 cells with AR-V7-mEGFP and (A-B) and AR-FL-mEGFP (C-D). Cells were transfected with AR-V7-mEGFP (A-B) or AR-FL-mEGFP (C-D) and were cultured in serum starved media. 48 hrs post transfection, foci were photobleached using confocal microscope with 40% of laser power for 10s. Time-lapse imaging on the bleached and reference foci were captured. (n=25 foci from 25 cells across three biological replicates). (E) PC3 cells were transfected with increasing amounts of either AR-FL-pcDNA or AR-V7-pcDNA plasmids in serum starved media. After 48 hrs, cells were stimulated with 1 nM DHT for 2 hrs. The cells were then harvested and the expression levels of AR-FL and AR-V7 in whole cell lysates were analyzed by western blots using AR-441 antibody that detects both AR-FL and AR-V7. (F) AR-V7-mEGFP and AR-FL-mEGFP condensates formation were visualized by confocal microscopy. (G) Foci density was quantified and plotted against expression levels. (H) Representative images of *in vitro* droplets formed by recombinant AR-FL-mEGFP-MBP-His or AR-V7-mEGFP-MBP-His in presence of 10% PEG and at each of the studied concentrations. Droplets were visualized by confocal microscopy and the number of droplets for each condition was recorded. (I) The number of droplets were quantified (n= three biological replicates and 15 images per replicate) and plotted as function of protein concentration.

AR-V7 and AR-FL are structurally different, with AR-V7 lacking the LBD. This structural difference may influence the ability of AR-V7 to form condensates. To assess whether AR-FL and AR-V7 exhibit similar phase behavior, we assessed whether they form foci at the same threshold concentration (C_threshold_). To do so, we transfected AR-negative PC3 cells with increasing concentrations of either AR-V7-mEGFP or AR-FL-mEGFP. Using identical expression plasmids, we observed higher expression levels of AR-V7 compared to AR-FL (Figure 3C). We quantified foci density for each isoform as a function of protein concentration, estimated using purified AR-FL protein as a reference (Figure 3C-G, S3A-B). AR-FL reaches a foci density of ∼2 at 0.7 μg/μl of expressed protein, whereas AR-V7 requires 1.5 μg/μl to reach a similar density, suggesting that AR-V7 has an approximately 2-fold higher concentration threshold (C_threshold_) than AR-FL. We next assessed the C_threshold_ *in vitro* using recombinant AR-V7-mEGFP and AR-FL-mEGFP tagged with a C-terminal Maltose Binding Protein (MBP) and a His-tag for purification purposes. We previously showed that MBP-His had minimal effect on AR phase behaviour *in vitro* ^18^. We verified protein concentrations using Coomassie-stained gels (Figure S3B) and then mixed increasing concentrations (2.5 nM to 60 nM) of each protein with 10% of the crowding agent polyethylene glycol (PEG 8000) to induce phase separation *in vitro* (Figure 3F-G). We found that AR-FL has an C_threshold_ of 20 nM +/-2 and AR-V7 an C_threshold_ of 30 nM +/-1, consistent with the higher C_threshold_ measured for foci formation in cancer cells. Taken together, our results are consistent with AR-V7 forming liquid-like, dynamic condensates in CRPC models but requiring ∼2-fold higher concentrations than AR-FL to do so.

### AR-V7 condensates require open chromatin for their formation

We previously demonstrated that AR-FL condensates colocalize with the enhancer of *FKBP5*, using DNA fluorescence in situ hybridization (DNA-FISH) ^18^. To further assess the role of chromatin accessibility and binding in the formation of AR-V7 and AR-FL condensates, we treated PCa cells with a PROTAC molecule, AU15330, which targets SMARCA2 and SMARCA4 proteins, the ATPase subunits of the SWI/SNF complex that plays a key role in chromatin remodeling ^46^. AU15330 has been shown to significantly reduce chromatin accessibility and the levels of euchromatin marker H3K27Ac ^46^. By analyzing ChIP-seq data from VCaP cells treated with AU15330 ^46^, we first confirmed that AU15330 reduces AR-FL recruitment to *FKBP5* enhancer sites in these cells (Figure S4A). We then treated 22Rv1 cells with 1 ¿M of AU15330 for 6 hrs, resulting in a significant reduction in SMARCA4 protein levels, as measured by western blot (Figure S4B). We also validated that AU15330 induced chromatin compaction, as demonstrated by the significant inhibition of chromatin digestion by MNase compared to untreated cells (Figure S4C) and by a marked increase in levels of the heterochromatin marker H3K9me2 (Figure 4A-B). Interestingly, we found that SMARCA4 also forms foci in 22Rv1 cells that colocalize with AR-V7-mEGFP condensates (Figure S4D), and that AU15330 reduced SMARCA4 foci, as measured by immunofluorescence (Figure 4A-B). Upon chromatin compaction, we found that condensate formation by both endogenous AR-V7 and AR-FL decreased (Figure 4A-B).

**Figure 4:**
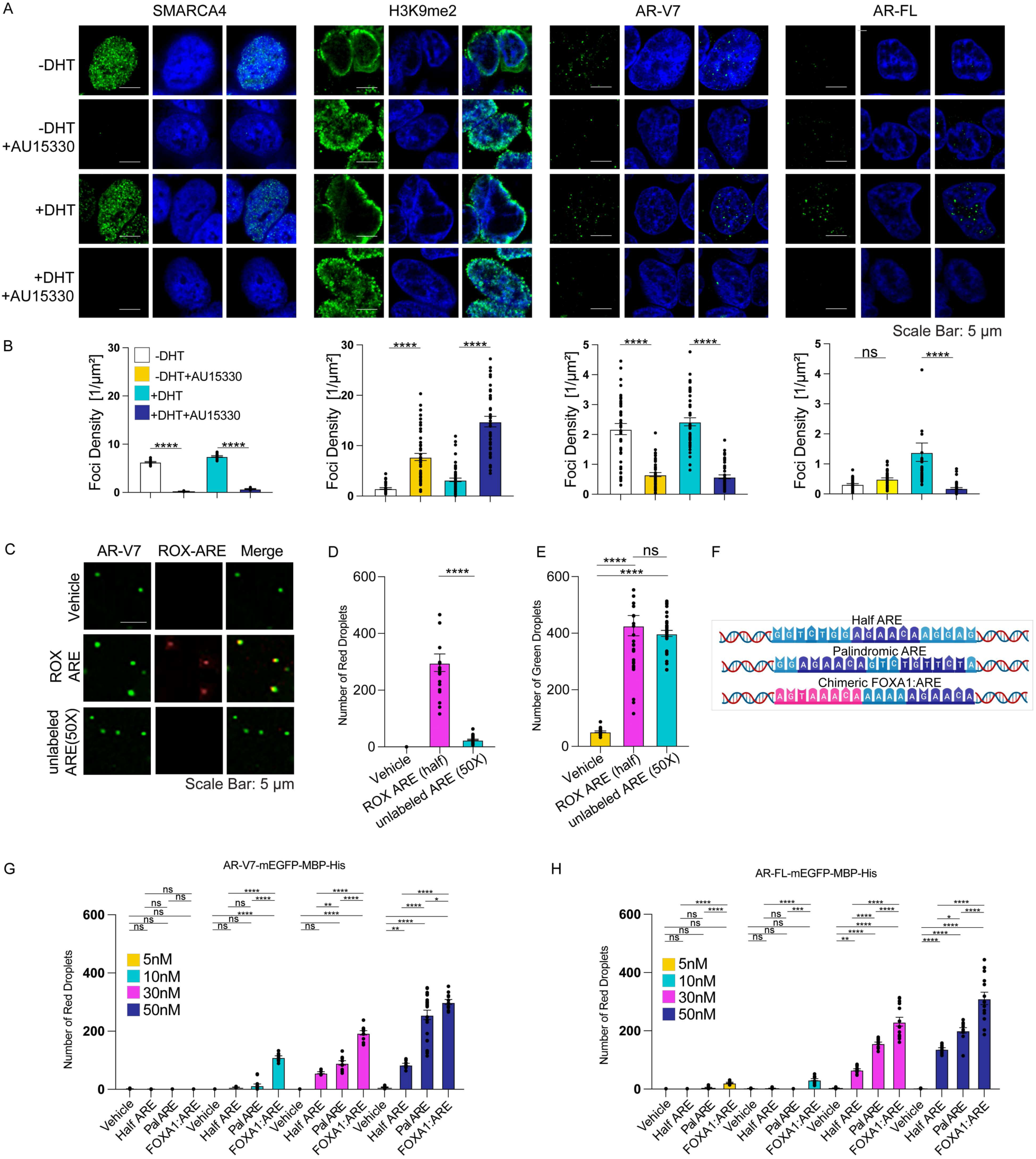
(A) 22Rv1 cells were serum starved for 48 hrs and then treated with vehicle or 1 nM DHT +/- AU15330 (1µM) for 6 hrs. The level of SMARCA4 and H3K9me2 as well as formation of condensates by endogenous AR-V7 and AR-FL were evaluated by immunofluorescence using confocal microscopy (n= three biological replicates and 15 images per replicate). (B) Quantification of foci density assessed by immunofluorescence. Data are presented as mean ± SEM. Statistical significance was determined using a t-test. (C) Recombinant AR-V7-mEGFP-MBP-His [50 nM] was used to form *in vitro* droplets in the presence of 10% PEG 8000, with either vehicle, ROX-labeled ARE [125 nM], or a 50X excess [6.25 μM] of unlabeled ARE. Samples were incubated at room temperature for 10 minutes before imaging with confocal microscopy. The numbers of red (D) and green (E) droplets were measured and plotted. Data are presented as mean ± SEM. Statistical significance was determined using a t-test. (F) The schematic of ARE sequences used in the *in vitro* assays. (G-H) Recombinant AR-V7-mEGFP-MBP-His or AR-FL-mEGFP-MBP-His was used at various concentrations to form droplets in 10% PEG 8000 with either vehicle or 125 nM ROX-labeled half-site, palindromic, or FOXA1:AR chimeric sequences. The number of red droplets was measured and plotted. Data are presented as mean ± SEM. Statistical significance was determined using a one-way ANOVA.

These results suggest that the accessibility of AR binding sites in euchromatin is crucial for AR-FL and AR-V7 condensate formation in 22Rv1 cells. Recently, Parolia *et al.* identified a chimeric AR binding motif4an FOXA1 element adjacent to an AR half site4as the most AR bound DNA sequence in LNCaP cells. This observation was supported by ChIP data from CRPC patients, showing greater enrichment of chimeric AR binding sites in the AR cistrome compared to samples from normal or primary PCa patients ^47^. Therefore, we tested how different DNA sequences (Figure S4D) affected droplet formation of AR *in vitro.* First, we assessed the colocalization of AR-V7 droplets with ROX-labeled ARE (half site 59-GGTCTGG**AGAACA**AGGAG**-39**) derived from a *KLK3* enhancer *in vitro*. We found that ROX-labeled ARE indeed colocalize with AR-V7 droplets (Figure 4C), and that this colocalization is reduced in the presence of a 50X excess of unlabeled ARE with the same sequence (Figure 4C-D). Additionally, we found that the presence of the ARE significantly increased the number of *in vitro* AR-V7 droplets by ∼8 times (Figure 4E). This aligns with our previous work and other studies showing that the presence of DNA lowers the threshold required for condensate formation ^18, 48^. Next, we investigated whether AR-FL and AR-V7 droplets showed a preference for specific DNA sequences and if these sequences affect their C_threshold_. We tested three ROX-labelled DNA sequences: palindromic **ARE** (59-GG**AGAACA**GTC**TGTTCT**A**-39**), a half-site **ARE** (59-GGTCTGG**AGAACA**AGGAG**-39**), and the chimeric ***FOXA1***/half-site **ARE** (59-AGTAAACA***AGTAAACA***AAAA**AGAACA**AGAACA**-39**) (Figure 4F), and examined their colocalization with AR-V7 and AR-FL droplets at varying protein concentrations (Figure 4G-H). For both AR-V7 (Figure S4E) and AR-FL (Figure S4F), all DNA constructs enhanced *in vitro* droplet formation compared to the vehicle. DNA constructs alone cannot form any droplets; thus, the red droplets represent DNA that is colocalized with AR droplets. Both AR-V7 (Figure 4G) and AR-FL (Figure 4H) showed the highest number of red droplets with the chimeric FOXA1-ARE sequence, followed by the palindromic and half-ARE constructs. Notably, the chimeric FOXA1-ARE sequence enables droplets formation at low AR concentrations, concentrations at which no droplets are observed for the other DNA sequences. Overall, these data emphasize the importance of chromatin and DNA accessibility for AR-V7 and AR-FL condensate formation and highlight the preferential colocalization of *in vitro* AR droplets with chimeric FOXA1-ARE sequences.

### AR-V7 condensate formation colocalize with transcriptional players

To evaluate whether AR-V7 condensates are transcriptionally active, we first examined their colocalization with phosphorylated MED1 (pMED1) and RNA polymerase II (pPol II) as do AR-FL condensates ^18^. We found that AR-V7-mEGFP condensates colocalize with pMED1 (Figure 5A) and pPol II in 22Rv1 (Figure 5B) cells in the presence and absence of androgens. To further validate these findings, we used proximity ligation assay (PLA) to detect proximity sites between pMED1 and pPol II, suggesting a potential interaction, and then confirmed colocalization of AR-V7-mEGFP with these interaction sites using fluorescence microscopy (Figure 5C). Consistent with our previous results for AR-FL, we also found that MED1-IDR lowers the C_threshold_ for *in vitro* droplet formation of AR-V7 ^18^. Specifically, AR-V7 droplets form at 12.5 nM in the absence of MED1-IDR and at 6 nM in the presence of MED1-IDR (125 nM) (Figure 5D-E). Our results demonstrate that AR-V7-mEGFP condensates associate with components of the transcriptional machinery in the 22Rv1 cells regardless of androgen stimulation.

**Figure 5.**
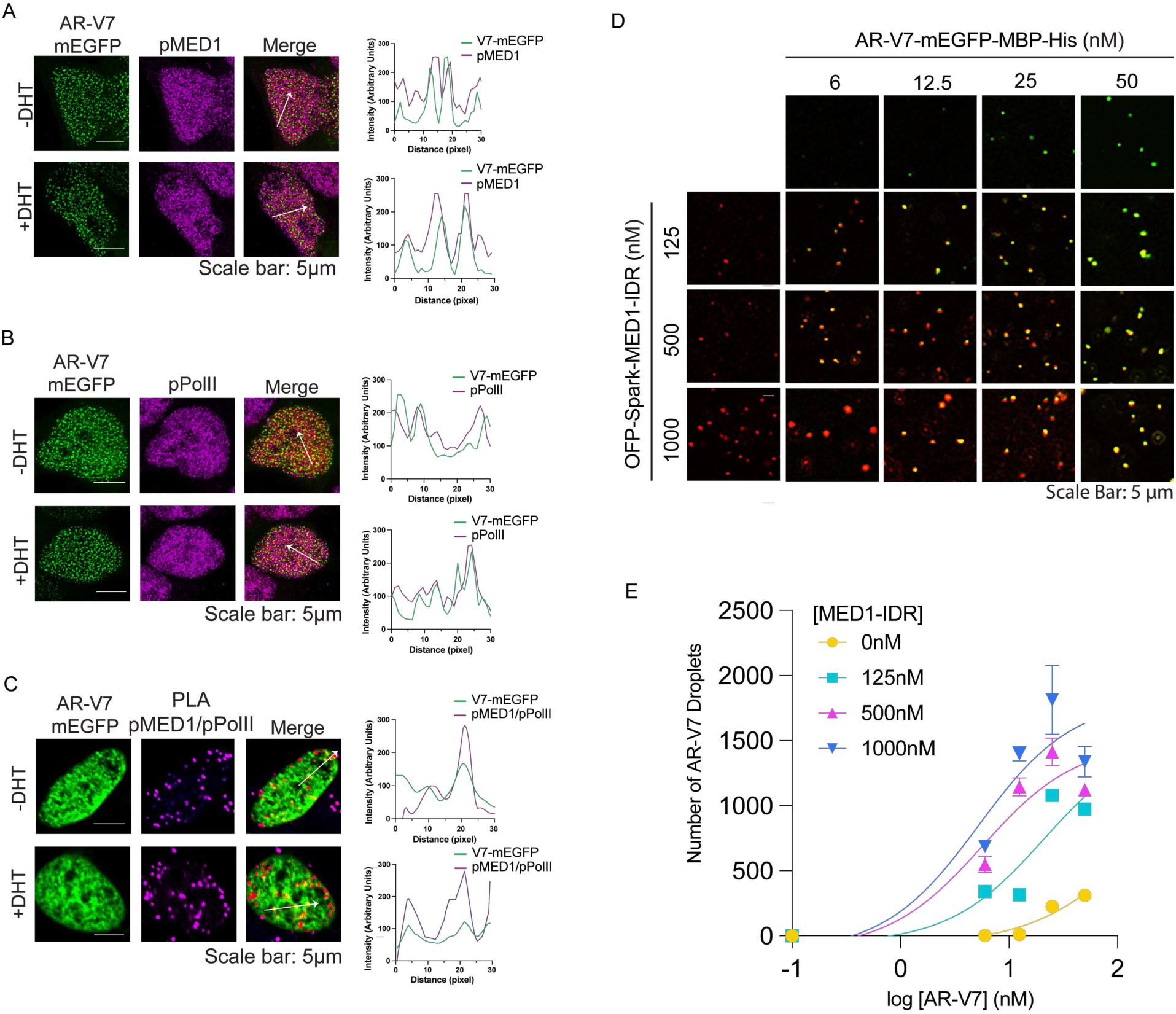
(A-B) 22Rv1 cells were transfected with AR-V7-mEGFP and cultured in serum starved media. 48 hrs following transfection, cells were stimulated with either 1 nM DHT or vehicle for 2 hrs and were then fixed. Endogenous pMED1 and pPol II were detected by immunofluorescence and their colocalization with AR-V7-mEGFP foci were evaluated by confocal microscopy. The original colors for pMED1 and pPol II signals were red. For compatibility with colorblind readers, the colors were adjusted in Photoshop using the Hue setting (red shifted by -65) (C) 22Rv1 cells were transfected with AR-V7-mEGFP and cultured in serum starved media. 48 hrs following transfection, cells were stimulated with either 1 nM DHT or vehicle for 2 hrs. The cells were then used in PLA for detecting the interaction of pMED1 and pPol II. Colocalization of AR-V7-mEGFP foci with pMED1 and pPol II interactions sites were evaluated by confocal microscopy. (n= three biological replicates and 15 images per replicate). (D) *in vitro* droplets of AR-V7-mEGFP-MBP-His were formed at different concentration and presence of various concentration of OFPSpark-MED1-IDR. The corresponding images for each condition are recorded. (E) The number of AR-V7 droplets (green) were quantified and graphed against MED1-IDR protein concentration.

### AR-V7 drives a condensate-dependent oncogenic program

We next wondered which transcriptional programs are dependent on the formation of AR-V7 condensates. We thus aimed to compare the transcriptomes of condensate-competent wild-type AR-V7 and a condensate-incompetent AR-V7 variant in AR-negative PC3 cells transiently transfected with these constructs. We used the Basu *et al.* ^20^ study to identify tyrosine (Y) residues to serine (S) mutants with reduced ability to form condensates. Specifically, we found that the 5YtoS mutant (Y481S/Y483S/Y504S/Y514S/Y531S) significantly reduced condensate formation in PC3 cells compared to the 4YtoS mutant (Y481S/Y483S/Y504S/Y514S) and wild-type AR-V7 (AR-V7-WT) (Figure 6A-C and S5A). We validated that the effect on condensate density of the 5YtoS mutant is not due to changes in expression levels (Figure 6D) nor nuclear translocation (Figure 6E). We also found that the AR-V7-5YtoS mutant reduces the expression of endogenous AR target genes (*FKBP5* and *TMPRSS2*) (Figure S5B) without affecting DNA binding as measured by ChIP-PCR (Figure S5C). Next, we used the 5YtoS mutant to identify transcriptomic programs that depend on AR-V7 condensates. We transfected PC3 cells with either AR-V7-WT-mEGFP or AR-V7-5YtoS-mEGFP for 48 hrs, isolated the RNA, and performed mRNA sequencing (RNA-seq). *FKBP5*, a shared target gene of AR-FL and AR-V7, showed the most substantial decrease in mRNA in the presence of the AR-V7 phase altering mutant (Figure 6F and Table S1). This finding aligns with our previous results, which showed that AR-FL condensates localize to the *FKBP5* gene enhancer, and that disruption of these condensates reduces *FKBP5* expression ^18^.

**Figure 6.**
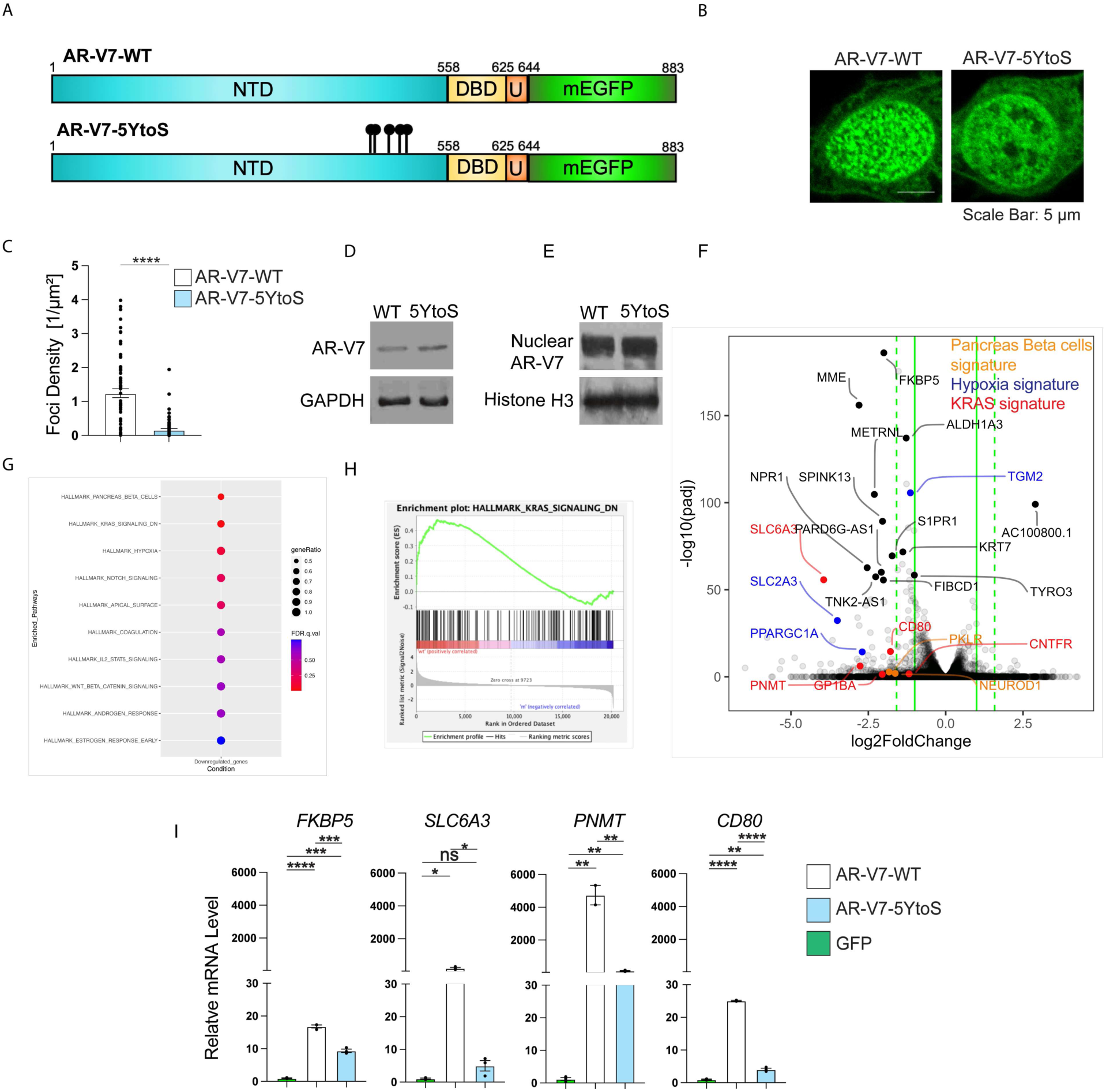
(A) Schematic of AR-V7-mEGFP-WT and AR-V7-mEGFP-5YtoS constructs. (B-C) PC3 cells were transfected with either wild-type or 5YtoS AR-V7-mEGFP (1.5 μg) and cultured in serum starved media for 48 hrs. The cells were then imaged by confocal microscopy and the condensate density of AR-V7-mEGFP-WT and AR-V7-mEGFP-5YtoS were quantified. Data are presented as mean ± SEM. Statistical significance was determined using a t-test. The corresponding whole cell lysates (D) as well as fractionated nuclear extracts (E) were evaluated by western blot. (F) PC3 cells were transfected with either AR-V7-mEGFP-WT or AR-V7-mEGFP-5YtoS and cultured in serum starved media for 48 hrs. RNA was then isolated for RNA-sequencing. The genes with fold change more than 2 and p-value less than 0.05 are shown in the scatter plot. (G-H) Pathway enrichment analysis by GSEA following RNA-seq of PC3 cells expressing wild-type or 5YtoS AR-V7-mEGFP. KRAS signaling pathway was one of the top enriched pathways. (I) RNA samples were isolated from PC3 cells transfected with either AR-V7-mEGFP-WT, AR-V7-mEGFP-5YtoS, or empty-mEGFP plasmid. Expression level of KRAS genes were detected by qPCR. Data are presented as mean ± SEM. Statistical significance was determined using a t-test.

We then performed a gene set enrichment analysis (GSEA) to identify enriched pathways and biological processes. The analysis highlighted the KRAS pathway as one of the most significantly enriched pathways (NES: 1.61, FDR q-value: 0.034) (Figure 6G-H). Consistent with this finding is a recent analysis of TCGA data ^49^, which demonstrated that PCa patients with high levels of AR-V7 exhibit significant enrichment in the KRAS pathway compared to those with low AR-V7 levels. To validate our finding, we examined the differential expression of *FKBP5* and the top altered genes in the KRAS signaling pathway (*SLC6A3*, *PNMT*, *CD80*) upon expression of AR-V7-WT and AR-V7-5YtoS, respectively, using qPCR. eGFP expression served as a control to confirm AR-V7-dependent regulation of these genes. Upon the expression of 5YtoS AR-V7 mutant in PC3 cells, we found a significant reduction in the expression levels *SLC6A3*, *PNMT*, and *CD80* (Figure 6I). As a control, we selected genes (*HNRNPA1*, *GRK2* and *PPP2R1B*) that were not affected by the disruption of AR-V7 condensates in our RNA-seq analysis, yet are known to be involved in PCa development and tumorigenesis ^50–52^. We did not find any significant differences in the expression levels of these genes between AR-V7-WT and AR-V7-5YtoS (Figure S5D). We thus identified a KRAS gene signature that is activated by AR-V7 condensates in PC3 cells.

### Validation of the KRAS signature in CRPC models

We next examined whether AR-V7-mediated expression of KRAS target genes occurs in AR-positive CRPC models, such as 22Rv1 and LN95, and whether these targets are specific to AR-V7 or shared with AR-FL. To this end, we assessed the expression of the identified genes in 22Rv1 and LN95 cells, which both express endogenous AR-FL and AR-V7. As AR-V7 is constitutively active, AR-V7 target genes should not see increased expression levels upon DHT stimulation, in contrast to genes under AR-FL control. We found that the expression levels of the identified KRAS gene signature did not change in the absence or presence of DHT (Figure S5A and S5B). In contrast, *FKBP5*, the target gene shared between AR-FL and AR-V7, shows increased expression upon DHT stimulation. To further demonstrate that the KRAS gene signature is independent of AR-FL, we measured gene expression levels following treatment of 22Rv1 and LN95 cells with the AR-FL PROTAC (ARV110; 100 nM) and in the presence of 1 nM DHT (Figure S5C and S5D). We observed no significant difference in the expression of the KRAS gene signature upon ARV110 treatment, while *FKBP5* displayed a marked reduction in its mRNA levels. These results suggest that the identified KRAS signature is AR-V7 specific in CRPC models. To demonstrate that this signature is dependent on condensate formation in these cells, we treated 22Rv1 (Figures S6E) and LN95 cells (Figure S5F), with either vehicle or 1% 1,6-hexanediol (1,6-HD) for 30 minutes. To rule out toxicity from 1,6-HD, we assessed cell viability and morphology, finding no significant changes (Figure S5E-F). Notably, the condensate density of endogenous AR-V7, detected by immunofluorescence, was reduced in both cell types after 1,6-HD treatment, which is associated with a significant decrease in KRAS gene expression, while control genes remain unaffected (Figure S6G-H). These results further validate that the identified KRAS genes are AR-V7-specific and that condensate formation is required for their transcription.

### AR-V7 binding site density determines condensate-dependent versus independent transcription

Surprisingly, our analysis of the AR-V7 condensate-dependent transcriptome did not identify known AR-V7 target genes such as *EDN2* ^53–57^ and *UBE2C*^38, 58–60^. We thus speculated that there may exist condensate dependent and condensate independent transcription mediated by AR-V7. There are multiple factors that can contribute to condensate-dependent and independent transcription including the binding site density ^61^, long-range chromatin interaction ^61, 62^ as well as epigenetic modifications ^63^. To explore this concept we focused on the AR binding site (ARBS) density as this could increase the local protein concentration and enhance condensate formation. Specifically, we examined whether the number of ARBS within the promoters (-20 kb) of affected genes (n=571) is higher compared to unaffected genes. For the unchanged genes, we randomly selected 571 genes from a compiled list of previously identified AR-V7 target genes (n=2335) for better comparison ^42, 49, 64, 65^ (Table S1). For both groups, we found that ARE half sites are the most abundant sites, followed by palindromic and chimeric FOXA1:ARE sites (Figure 7A). Although the number of binding sites for chimeric FOXA1:AR and full ARE is small, the changes in their abundance are significant, while the half AREs exhibit a larger number of binding sites and a more pronounced change. It has been proposed that condensate formation of transcription factors could enable prolonged and robust transcription bursts ^66^. Therefore, we transfected PC3 cells with varying concentrations of AR-V7-mEGFP and measured the expression of genes affected by the loss of condensate formation (*SLC6A3, PNMT, CD80*) as well as known AR-V7 target genes (*EDN2*^53–57^*, UBE2C*^38, 58–60^) whose expression was unaffected by the loss of condensate formation (Figure 7B). qPCR results revealed an exponential increase in gene expression with increasing AR-V7 levels for condensate-dependent genes, whereas condensate-independent target genes show a much lower and linearly increasing level of expression as a function of increasing AR-V7 concentrations (Figure 7B-C). In summary, AR-V7 mediates both condensate-dependent and independent transcription, with condensate-dependent genes showing a pronounced increase in expression at higher AR-V7 levels, while condensate-independent genes exhibit a more gradual increase. Our results suggest that the increased number of binding sites could contribute to such change in transcriptional output.

**Figure 7.**
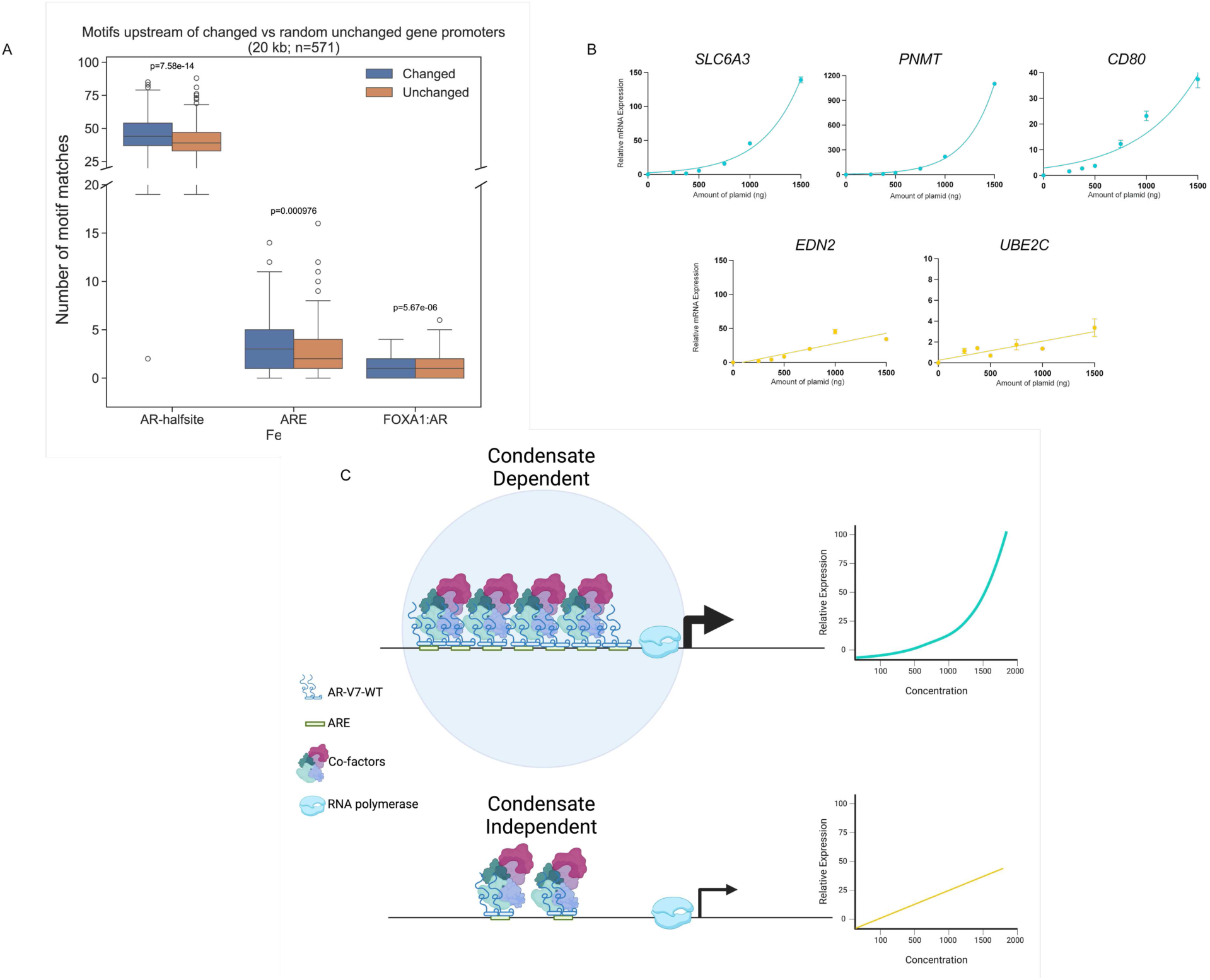
(A) The number of half sites, palindromic sites, and chimeric FOXA1:ARE sites in the promoters (-20 kb) of genes with altered expression in the AR-V7-5YtoS mutant compared to AR-V7-WT (blue, n∼500) and an equivalent number of unaffected genes (orange, n∼500). (B) PC3 cells were transfected with varying concentrations of the AR-V7-mEGFP expression plasmid, and the expression of target genes was quantified relative to PC3 cells transfected with the GFP plasmid. (C) Schematic illustrating two paradigms of transcription: condensate-dependent and condensate-independent. The top panel depicts genes with multiple AR-V7 binding sites, leading to increased accumulation of AR-V7 and its co-factors, resulting in exponential target gene expression. The bottom panel shows condensate-independent transcription, where a limited number of binding sites leads to lower and more linear target gene expression.

## Discussion

Dysregulation of biomolecular condensates has been associated with various cancers such as prostate, lung, breast, pancreatic, ovarian, and colorectal cancer. In many cancers, oncogenic drivers form condensates at specific genomic regions to orchestrate the transcription of programs required for cancer cell survival, growth and metastasis ^3^. In advanced PCa and specifically CRPC, the expression of constitutively active AR-V7 is one of the mechanisms that has been linked to aggressive tumor behaviour and treatment resistance ^67^. In this study, we found that endogenous as well as exogenously expressed AR-V7 forms nuclear foci independently of androgens in several CRPC models. Previously, we ^18^ and others ^22, 62^ found that exogenously expressed AR-V7 either does not or minimally forms foci in LNCaP cells, which are androgen-sensitive cells not expressing any AR splice variants. In contrast, LN95 cells, adapted to grow in castration conditions, display significant foci density when AR-V7 is expressed in these cells at similar level as in LNCaP cells. Our data do not support the presence of cytoplasmic foci observed by Thiyagarajan *et al.* ^33^ when they exogenously expressed AR-V7 in kidney COS7 cells, suggesting that their finding may be specific to the cell model they used. Importantly, we demonstrate that AR-V7 forms foci in CRPC models independently of AR-FL. Although, prior studies suggested that AR-V7 can heterodimerize with AR-FL and help the latter to translocate to the nucleus and activate transcription in the absence of androgens^21, 33, 41^, we did not find any evidence for AR-V7-assisted nuclear translocation or condensate formation by AR-FL.

Our data support the idea that the foci formed by AR-V7 in CRPC models, similar to those formed by AR-FL in various PCa models, represent biomolecular condensates. AR-V7 foci display diffuse exchanges that are similar to those previously measured for foci formed by AR-FL ^18^. Interestingly, we found that AR-V7 requires about twice the expression level in cells and in vitro to form condensates compared to AR-FL. However, expression levels alone are not sufficient, as AR-V7 forms condensates in LN95 cells but not in LNCaP cells, despite similar expression levels. This raises the question of why AR-V7 can form condensates in some CRPC models but not others. It is likely that cell model- or tumor-stage-specific changes in key transcriptional coregulators impact AR-V7 condensate formation. Indeed, similar to AR-FL ^18^, we found that co-activators like MED1-IDR can lower the C_threshold_ for AR-V7 condensate formation *in vitro*. Moreover, we found that the chimeric FOXA1-ARE sequence lowered the C_threshold_ for both AR-FL and AR-V7 droplets by two to three-fold while ARE-half did not have drastic effect. This result is notable, given the previous finding that AR-bound enhancer regions exhibit a higher prevalence of chimeric FOXA1:ARE in CRPC when compared to primary PCa or normal tissues^68^. Thus, cell-model specific changes in the presence or accessibility of specific AREs are also likely to impact condensate formation by AR-V7.

It has been established that AR-V7 contributes to the aggressive PCa behaviour and treatment resistance ^67^. However, it remained largely elusive how this AR variant drives specific oncogenic programs. Using a mutational variant (AR-V7-5YtoS) that reduces condensate formation in CRPC models, we revealed that the genes most affected in their expression by the altered phase behaviour of AR-V7 are the same genes that are elevated in CRPC and linked to treatment resistance such as *FKBP5* ^53^*, ALDH1A3* ^69^*, SPINK13* ^70^, and *TYRO3* ^71, 72^. Moreover, a GSEA analysis of genes susceptible to changes in AR-V79s ability to form condensates identified KRAS signaling members as particularly affected. Follow-up experiments in 22Rv1 and LN95 confirmed that the expression of the identified genes from the KRAS signaling signature follows changes in AR-V7 condensate formation (Figures S6). These findings highlight the importance of AR-V7 condensates in regulating PCa-associated programs, particularly with respect to members of the KRAS signaling pathway. KRAS has been identified as a transformative factor that enhances prostate cancer stemness and bone metastasis via CD24 upregulation ^37^. Additionally, the MYC-associated zinc-finger protein (MAZ) has been shown to induce bone metastasis in prostate cancer through KRAS signaling ^73^, and a KRAS fusion with the UBE2L3 gene was observed in DU145 CRPC cells that metastasized to the brain, promoting cell invasion ^36^. Expressing this fusion in normal prostate epithelial cells resulted in increased cellular proliferation and invasion^36^.

Surprisingly, we found that known AR-V7 target genes, such as *UBE2C* ^38, 58–60^*, EDN2* ^53–57^*, SLC3A2* ^49^, and *ETS2* ^54–56, 64^, are not showing differential expression when using AR-V7-5YtoS, suggesting that the expression of these genes is not modulated by condensate formation. By analyzing the number of AR binding sites in condensate-independent genes previously identified as AR-V7 targets and comparing them to condensate-dependent genes, we observed a significant increase in binding sites in genes affected by the loss of condensate formation. Notably, condensate-dependent genes exhibited an exponential increase in expression compared to the linear expression of condensate-independent genes. Such a <boosted= expression has been associated with genes that are regulated by transcription factors able to form transcriptional condensates ^3^. Our findings reveal two paradigms of transcription mediated by AR-V7: condensate-dependent and condensate-independent transcription (Figure 7C). A higher number of binding sites recruits more AR-V7 proteins, which increases the local AR-V7 concentration and potentially facilitates condensate assembly and boosts gene expression under the condensate-dependent regime. Such mechanism may particularly benefit genes for which the transcription factor has lower affinities. Condensate formation could ensure a longer dwelling time of the transcription factor on chromatin, thereby prolonging and enhancing the expression of these genes^74, 75^.

## Material and methods

### Resources and Materials

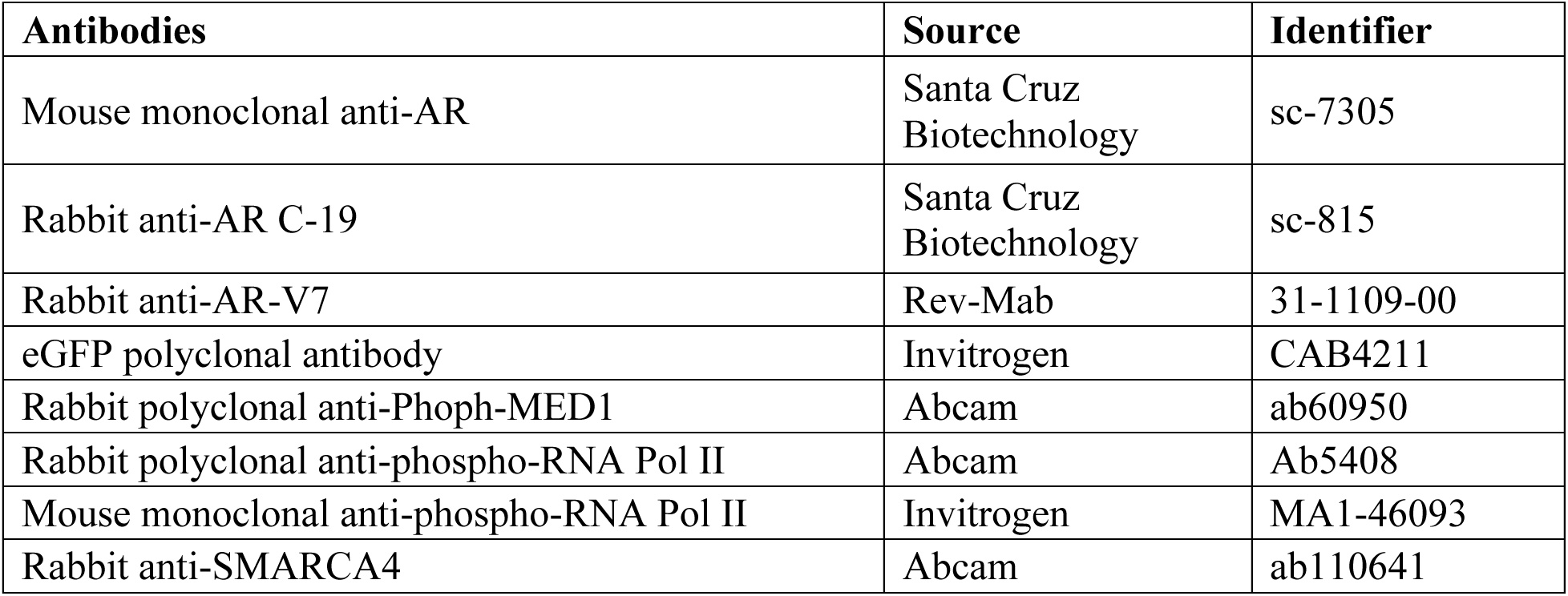
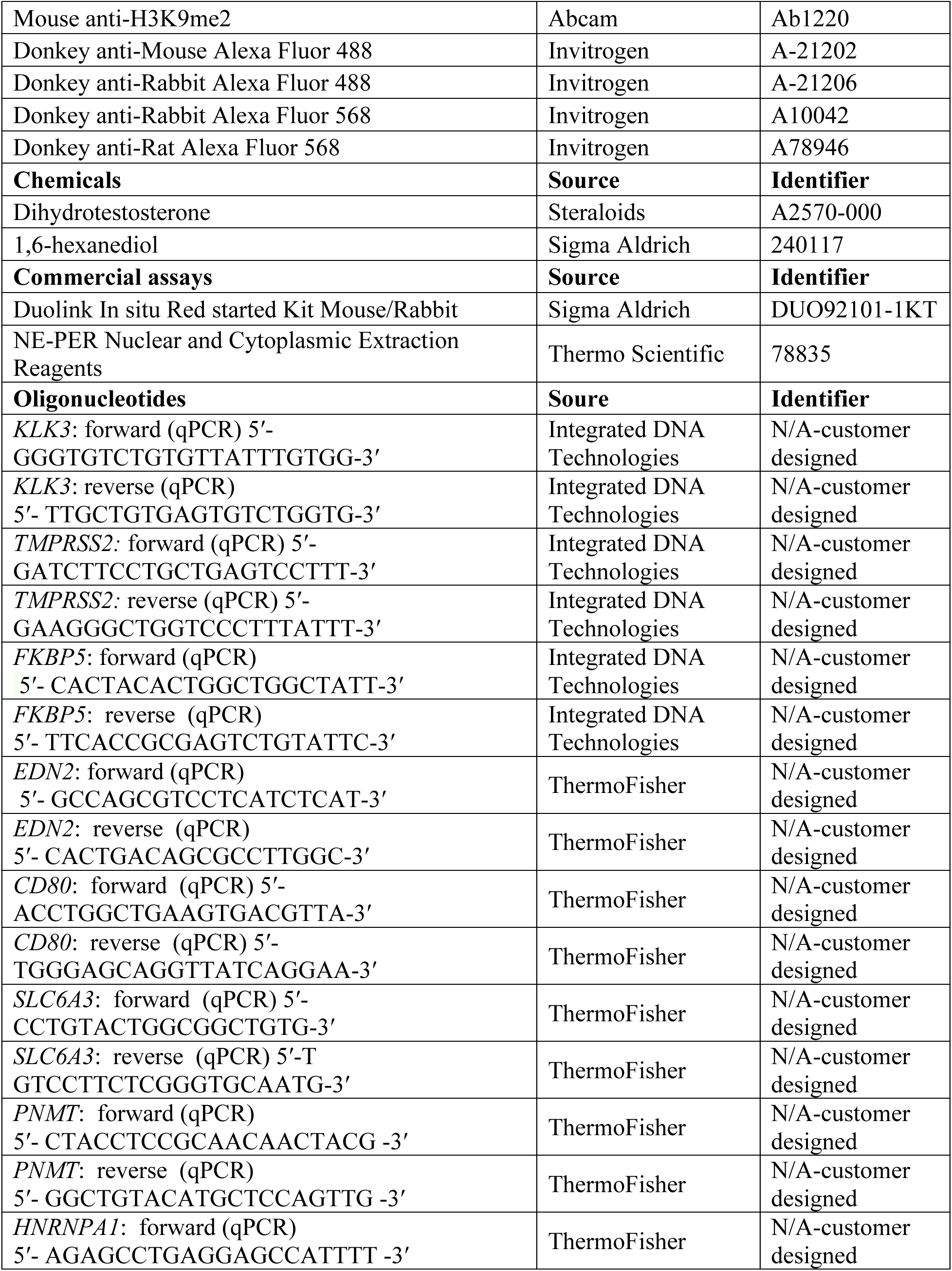
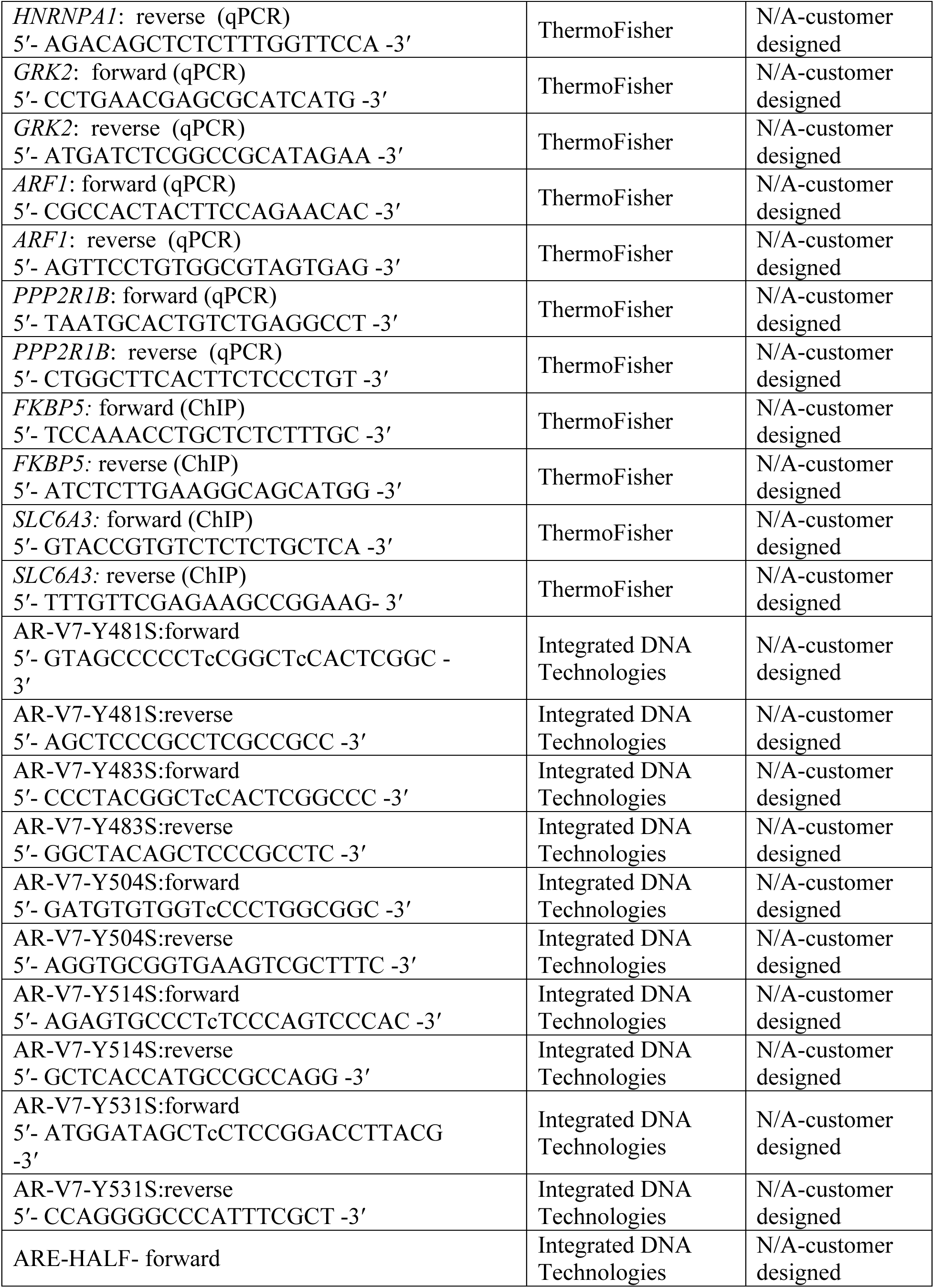
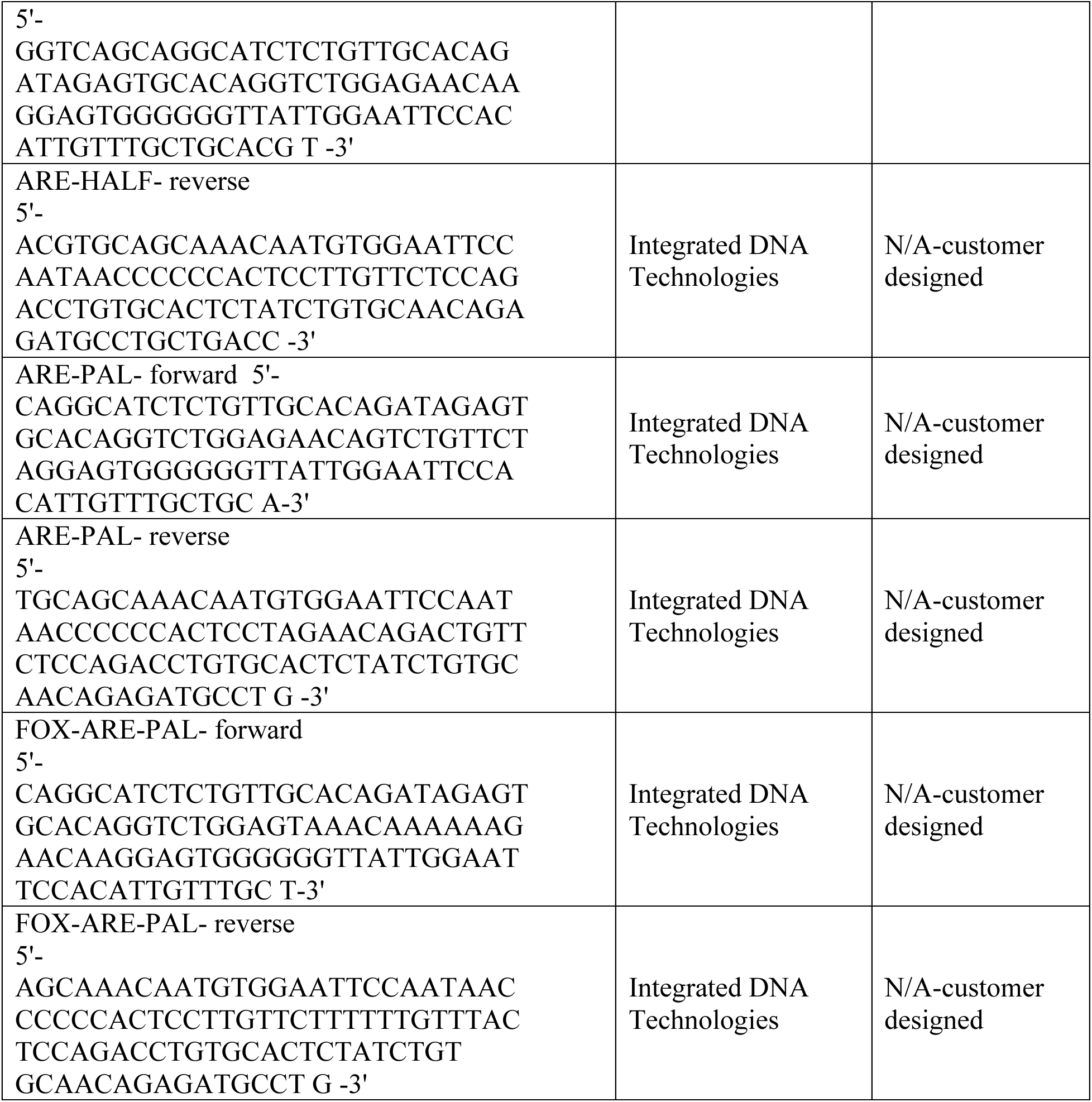

### Cell lines

LNCaP, LN95, 22Rv1, VCaP, HEK293t cells were received from the American Type Culture Collection (ATCC). LNCaP, 22Rv1 and HEK293t cells were grown in RPMI-1640 media containing 10% fetal bovine serum (FBS). LN95 cells were grown in phenol red free RPMI-1640 containing 10% charcoal strip serum (CSS). VCaP cells were grown in DMEM containing 10% FBS. Cell lines were authenticated by STR profiling.

### Transfection

Transfection of plasmids were carried out using TransIT2002 (Mirus) using the manufacture guideline (Mirus Bio-MIR5404)

### Immunofluorescence staining and confocal microscopy

Cells grown on glass coverslips were fixed with 3:1 Acetone: Methanol mixture for 15 min. Cells were then washed with ris-Buffered Saline with Tween 20 (TBST) and permeabilized with 0.5% Triton X-100 for 5 min at room temperature. Cells were then washed with TBST and blocked with 4% Bovine Serum Albumin (BSA) in Phosphate-Buffered Saline (BPS) solution for 1 hr at 37 çC. The cells were then incubated with antibody diluted in 4% BSA in PBS at 4 çC for overnight. Cells were then washed with TBST and incubated with the Alexa-Flour secondary antibodies (1:500) at 37 çC for 1 hr in the dark. Coverslips were then mounted on slides with DAPI containing mounting medium (VECTOR).

### Confocal microscopy and quantification

Fluorescence imaging was performed using Olympus FV3000RS confocal microscope (Olympus). Images were taken using the FV31S-SW software. The 60x UPLAPO oil objective and Galvano scanner with scanning at 2 ¿sec/pixel were used to capture images. The confocal pinhole was set to automatic and airy disk was set to 1.00. The 405 nm, 488 nm and 564 nm lasers were used. Gain and offset were set to 1.00 and 3.00 respectively and the power and voltage were set to the same condition within each experiment.

Image processing and quantification was performed using the CellSense software (Olympus) with a customized in-house Macro script with the following steps: 1) Select DAPI nuclei, 2) Set DAPI nuclei as regions of interest (ROI), 3) Select GFP/RFP detected ROI and quantify Foci. Setting up a macro was performed on 10 sample images and were confirmed its accuracy on calling foci in subsequent images. To select DAPI nuclei, first the DAPI channel (405 nm) was selected, then using automatic threshold, count and measure function was performed. To set the DAPI nuclei as ROI, first all DAPI positive regions were selected, then ROIs were created from these selected objects. After the ROIs are established, the objects were then deselected. The last step is detecting the GFP/RFP positive cells to perform quantification. To do so, GFP or RFP channels were selected. Using manual threshold, count and measure function was performed on the GFP/RFP channels on ROIs. This threshold was set to ensure selection of all GFP-RFP positive cells. Now all the detected objects were selected and new ROIs were created using the GFP/RFP positive objects. Upon setting the new ROIs, the selected objects were deselected. After selection of all GFP/RFP positive cells, a count and measure function is performed to record the total intensity of fluorescence in the detected ROIs. Then using Adaptive threshold, count and measure on ROI function was performed on 488 or 564 channels. This threshold was set to insure detecting all foci in the nucleus. The resulting measurements which includes number of foci, ROI area and object density were recorded. The macro were ran for multiple samples in each experiments to make sure the detection threshold is optimized.

### Proximity Ligation in situ Assay (PLA)

Cells grown on coverslips were washed with PBS and fixed with 3:1 Acetone:Methanol mixture for 15 min at room temperature, followed by permeabilization with 0.5% Triton-X-100 in PBS for 5 min with shaking. Cells were blocked using solution provided by the Duolink In Situ Detection Reagent Fluorescence Kit (Sigma-Aldrich, Cat# DUO92008) for 1 hr at 37çC and incubated with antibody mixture (anti-phospho-MED1 and anti-phospho-Pol II) overnight at 4çC in a humidified chamber. The slides were then incubated with anti-rabbit Plus and anti-mouse Minus PLA probes conjugated to specific oligonucleotides for 1 hr at 37çC. Next, the oligonucleotides were hybridized and ligated for 30 min at 37çC and amplified for 120 min at 37çC using ligase and polymerase provided in the kit. The slides were then visualized using Olympus Confocal Laser Scanning Microscope.

### Quantitative real-time PCR

RNA was extracted from cells using TRIzol (Thermo Fisher, Cat# 15596026) according to manufacture9s instruction. For cDNA synthesis, 2 μg of total RNA was used and converted to complementary DNA using cDNA synthesis kit (Thermo Fisher, Cat#. E3010). The sequences of the primers used for PCR amplification are detailed in mateirals and methods. SYBR Green PCR master mix (Roche Diagnostics, Cat# 4913850001) was used for amplification and analyzed in ABI QuantStudio 7 pro. Fast-Real-Time System (Applied Biosystems). The PCR cycles were: 2min at 95çC, followed by 10 min at 95çC, then 40 cycles of 95çC for 15 sec and 60çC for 1 min. Relative fold changes were calculated using 18s gene as an internal control.

### Fluorescence Recovery After Photobleaching (FRAP)

FRAP analysis was performed using Olympus *FVS3000 confocal microscope with 60XUPLAPO objective and oil immersion*. The selected condensates were bleached with 10% of laser power, and the fluorescence recovery was measured for 30sec post bleaching. On the same filed, a reference non-bleached condensate was captured simultaneously. Half-lives and mobile fractions were analysis using CellSense software (Olympus) and graphed using GraphPad Prism 9. A total of 25 condensates in 3 biological replicates were monitored for each cell line.

### Western blot

The extract of cells were prepared using 1% NP-40 lysis buffer along with 5mM NaCl, 1mM orthovanadate, 10mM iodoacetamide, 1mM EDTA, 0.25% Na Deoxycholate, 1mM Phenylmethylsulfonyl Fluoride (PMSF) and 1X protease inhibitor cocktail (Roche Diagnostics). For SDS-PAGE, 50-100 μg of cell lysate were denatured and separated using 10% Tris-glycine gels then transfered to PVDF membrane. Membranes were them blotted with denoted antibodies and detected with ECL (Pierce-ThemoFisher Cat#32106).

### Viability assay

Cells were cultured in 96-well plates with a 100 µl culture volume per well, reaching 80% confluency 24 hrs after seeding. Afterward, the cells were treated with either vehicle or 1% Hexanediol for 30 minutes. Subsequently, 100 µl of 2X CyQUANT detection reagent was added to each well, and the plates were incubated for 60 minutes at 37°C. Fluorescence was measured using excitation at 480 nm and emission at 535 nm.

### Expression and purification

AR-V7-mEGFP-MBP-His or the AR-FL-mEGFP-MBP-His) was cloned in pSF-CMV-Puro-COOH-TEV-GST plasmid (Sigma-Aldrich). This construct was then transfected at 1mg/ ml DNA and 3 mg/ml of Polyethylenimine (PEI 25K) in FreeStyle 293-F cells (Gibco Cat#. R790-07) with 0.5 x 106 cells/ml cells density and >95% cells viability. Cells were maintained at 37 çC with 8% CO2 while shaking at 90 rpm. 48 hrs post transfection, cells were harvested and resuspended in buffer A (50 mM Tris, pH 8.0; 150 mM NaCl; 5% glycerol, 20 mM ZnSO4; 0.5% NP40; 0.2 mM TCEP; and 0.1 mM PMSF, 20 mM DHT for AR-FL) supplemented with 0.1 mg/ml DNaseA, 5 mM MgCl2, complete protease inhibitors (Roche, 11873580001) and 1 mM PMSF. The cells were lysed using dounce (40 times) and following centrifugation, AR-V7-mEGFP-MBP-His or AR-FL-mEGFP-MBP-His were purified by affinity chromatography using amylose agarose resin. All the fractions were subjected to gel analysis using 10% SDS-PAGE and confirmed the protein of interest by Western blot and mass spectrometry. Fractions containing protein of interest were pooled and concentrated (using Amicon Ultra centrifugal filters, 100K MWCO). For AR-FL-mEGFP-MBP-His, concentrated fractions were injected on size exclusion chromatography (Superose 6) and pure non-aggregated fractions pooled, concentrated and used to study the *in vitro* droplet formation.

### *In-vitro* droplet assay

Purified AR-V7-mEGFP-MBP-His protein or AR-FL-mEGFP-MBP-His were diluted in the assay buffer (50mM Tris, pH 8.0; 150 mM NaCl; 5% glycerol; 20 μM ZnSO_4_; 0.05% NP-40; 0.2 mM TCEP; and 0.1 mM PMSF) and was mixed with 1 or 10 μM of DMSO or inhibitors. The mixture was then mixed with 10% crowding agent (PEG-8000). Next, 10 μl of sample mixture was loaded on glass bottom 35mm petri dish (MatTek, Cat# P35G-0-14-C) and incubated for 10min. The samples were imaged with Olympus FVS3000 confocal microscope with 60XUPLAPO objective and oil immersion. For colocalization of MED1 with AR-V7 droplets, purified MED1-IDR-OFPSpark (residues 948-1574) were mixed at specified concentrations with AR-V7-MEGFP-MBP-His, in presence of 10% PEG-8000. The mixture was then added to the glass bottom petri dish for visualization following 10min incubation at room temperature.

### Library preparation and RNA sequencing

Total RNA integrity is assessed with the TapeStation 4200 (Agilent Technologies; RNA Assay Cat No.5067-5576) and quantified with the Qubit RNA HS assay (ThermoFisher; Cat No. Q32852). 200 ng of total RNA is used as input to enrich for mRNA with the NEBNext Poly(A) mRNA Magnetic Isolation Module (NEB; Cat No. E7490). Strand-specific libraries are prepared using the MGIEasy RNA Directional Library Prep kit (MGI; Cat No. 1000006386). Library quality and size are checked with the TapeStation 4200 (Agilent Technologies; D1000 Assay Cat No. 5067-5582) and then quantified using the Qubit dsDNA HS Assay (ThermoFisher; Cat No. Q32854). Single stranded circular libraries and DNA nanoballs (DNBs) were prepared following manufacturer’s instructions and sequenced using a Complete Genomics DNBSEQ-G400RS High-throughput Sequencer to generate about 20 million 100-bp paired-end reads per library. Fastq files are aligned to the reference transcriptome GRCh38_108 with STAR (version 2.6.0c), quantified with HTSEQ (version 0.11.2) and normalized/differential expression calculated with DESeq2 (version1.16.1).

### RNA-seq data analysis

Gene Set Enrichment Analysis (GSEA) (https://www.gsea-msigdb.org/gsea/index.jsp) was performed to identify significantly enriched pathways in PC3 cells transfected with wild-type AR-V7-mEGFP compared to those transfected with 5YtoS AR-V7-mEGFP.To do so, expression data sets were prepared, were count reads lower than 10 were eliminated from samples. Following parameters were for analysis; Gene sets database: h.all.v2024.1.Hs.symbols.gmt [Hallmarks], Number of permutations: 1000, Collapse to gene symbols, Permutation type: gene set and Chip platform: Human_Ensemble_Gene_ID_MSigDB.v2024.1.Hs.chip.

### Quantification of AR binding sites

GENCODE v19 genes were used for analysis. 20 kb sequences upstream of the transcripts of condensate dependent genes were obtained using bedtools (<flank= and <getfasta=). These sequences were scanned for the half ARE, full ARE and FOXA1:AR composite motifs available in the HOMER database using HOMER (annotatePeaks.pl -size given -nmotifs). To compare the abundance of motifs to condensate independent genes (n=571), we compiled a set of ARv7 regulated genes from other studies ^42, 49, 64, 65^, subtracted the condensate dependent genes (n=2335). From this set of condensate independent but ARv7 regulated genes, we randomly selected 571 for better comparison. Mann Whitney U test was performed between the changed and unchanged groups for statistical analysis. Visualization was performed using seaborn.

### Chromatin Immunoprecipitation

One 15 cm plate of PC3 cells (∼10 x 10^6^ cells) per immunoprecipitation was transfected with AR-FL-mEGFP, AR-V7-mEGFP, or AR-V7-5YtoS-mEGFP in starved cultured media. 48 hrs following transfection, the AR-FL transfected cells were stimulated with and without 1 nM DHT for 2 hrs. All the cells were crosslinked with 15ml of 1% formaldehyde in phenol red-free media for 10 min at room temperature and quenched with 1.5ml of 1M glycine. The cells were then washed twice with cold PBS and harvested in PBS containing protease inhibitor. Cells were then pelleted and resuspended in lysis buffer 1, rotated at 4°C for 10 min and centrifuged. The pellet was then resuspended in lysis buffer 2 and rotated at 4°C for 5 min and centrifuged. Following centrifugation, the pellet was resuspended in 500µl of lysis buffer 3. The lysate was then sonicated at an amplitude of 10 for 30 seconds on and 30 seconds off for a total of 5 min and run on an agarose gel to confirm sufficient sonication. The lysate was then diluted to 1ml with lysis buffer 3 including Triton X-100 to a final concentration of 1%. 50µl of input sample was removed and remaining lysate was added to the antibody bound beads following three washes in blocking buffer and rotated overnight at 4°C. For antibody preparation, 50µl of dynabeads per immunoprecipitation were washed in blocking buffer (PBS, 0.5% bovine serum albumin) and mixed with 10µg of antibody: AR 441 and normal mouse IgG in 500µl of blocking buffer overnight at 4°C. The next day, the beads were washed 6 times with 1ml RIPA buffer and once with 1ml TE. 200µl of elution buffer was added and vortexed. ChIP and input samples were incubated at 65°C overnight to reverse the cross-linking. The following day, the supernatant was separated from the beads and 200µl of TE was added to all samples. 8µl of RNase A (1mg/ml) was added and incubated at 37°C for 30 min. 4µl of Proteinase K (1mg/ml) was added and incubated at 55°C for 1 hour. DNA was isolated with the DNA cleanup kit (Qiagen) and evaluated with qPCR.

### Statistics

Statistical analysis were performed using two-tailed unpaired Student9s *t*-test. *P* values are indicated by start were ns ≥ 0.05, * 0.01 to 0.05, ** 0.001 to 0.01, *** 0.0001 to 0.001, **** < 0.0001.

## Acknowledgement

S.M. is the recipient of a Michael Smith Health Research BC 2020 Research Trainee Award [RT-202030455]; N.P and J.F. is the recipient of 2023 and 2021 Prostate Cancer Foundation BC Grant-in-Aid award respectively; J.G. is supported by the Canadian Institutes of Health Research [AWD-017620]; NSERC. The project described was also supported by award number P50 CA097186 from the National Cancer Institute. Funding for open access charge: start-up fund for Dr Lallous as Early career investigator.

## Contribution

Conceptualization: NL, JG. Methodology: S.M., N.P., J.F. Validation: S.M., N.P., J.F., Formal Analysis: S.M., J.F., N.P., T.E.K., N.A.L., N.L., R.B., S.K.R., S.V., J.M.B. Investigation: S.M., J.F., N.P., S.D., M.G., A.H., M.T., Resources: N.L., M.E.G, C.C.C, Data Curation: R.B., S.M., T.E.K., S.L.B. Writing-Original Draft: S.M., N.L. Writing-Review & Editing: All authors. Visualization: S.M, R.B., T.E.K., S.K.R. Supervision: N.L., Project Administration: N.L., S.M., Funding Acquisition: N.L., J.G., S.M.

## Supplemental Figure Legends

**Supplementary Figure S1.**
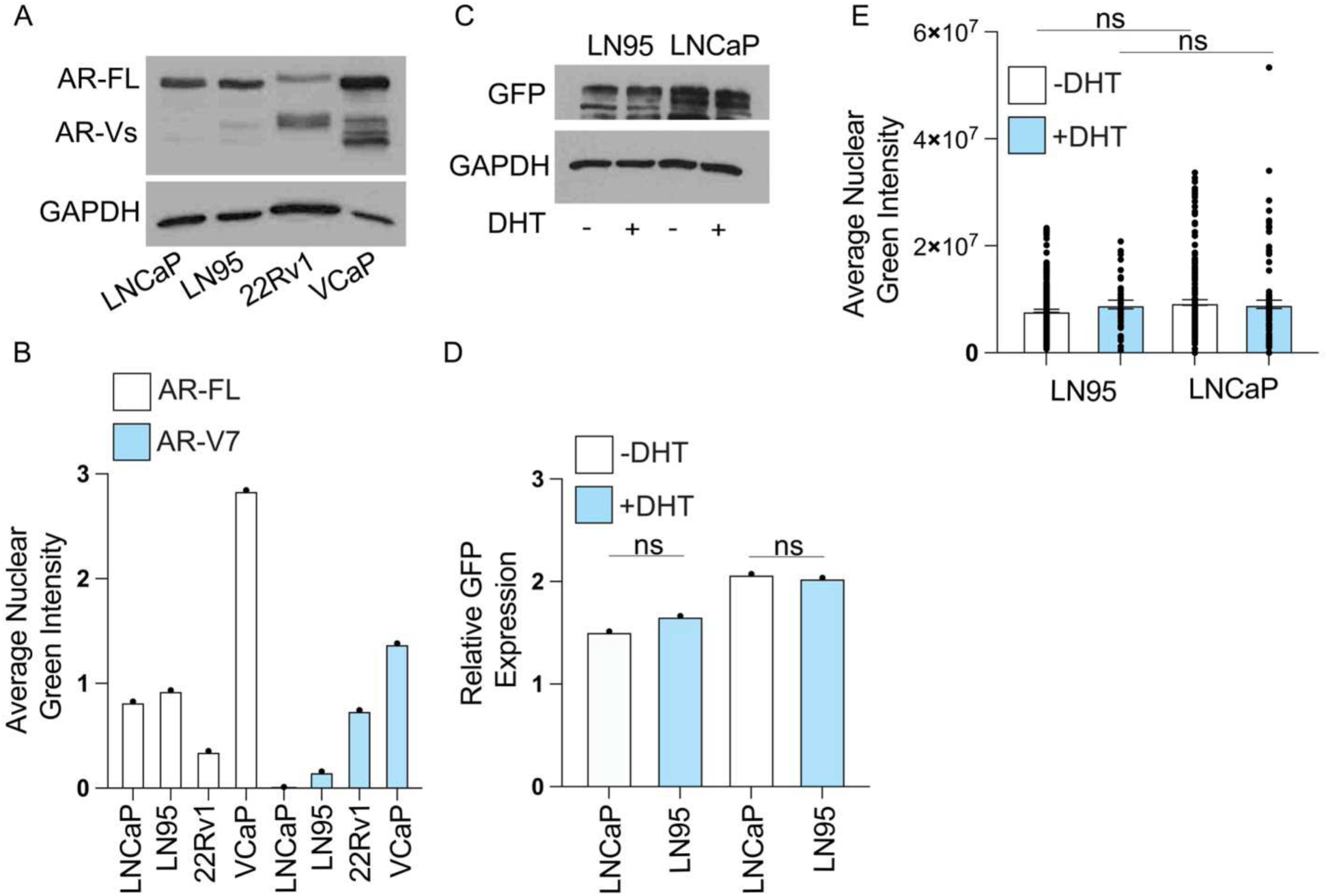
(A) Western blot showing the expression levels of endogenous AR-FL and AR-V7 in the studied PCa models using AR-441 antibody targeting N-terminus. (B) Densitometric quantification of AR-FL and AR-V7 western blot bands normalized to GAPDH. (C) LNCaP and LN95 cells were transfected with AR-V7-mEGFP in serum-starved media. 48 hrs post-transfection, cells were stimulated with either vehicle or 1 nM DHT. Cells were then lysed, and AR-V7-mEGFP expression levels were detected using a GFP antibody. (D) Densitometric quantification of AR-V7-mEGFP expression in LNCaP and LN95 cells. (E) LNCaP and LN95 cells transfected with AR-V7-mEGFP were stimulated with 1 nM DHT for 2 hrs. Cells were fixed and imaged via confocal microscopy. The graph shows the average nuclear green intensity per cell. Data are presented as mean ± SEM. Statistical significance was determined using a t-test.

**Supplementary Figure S2:**
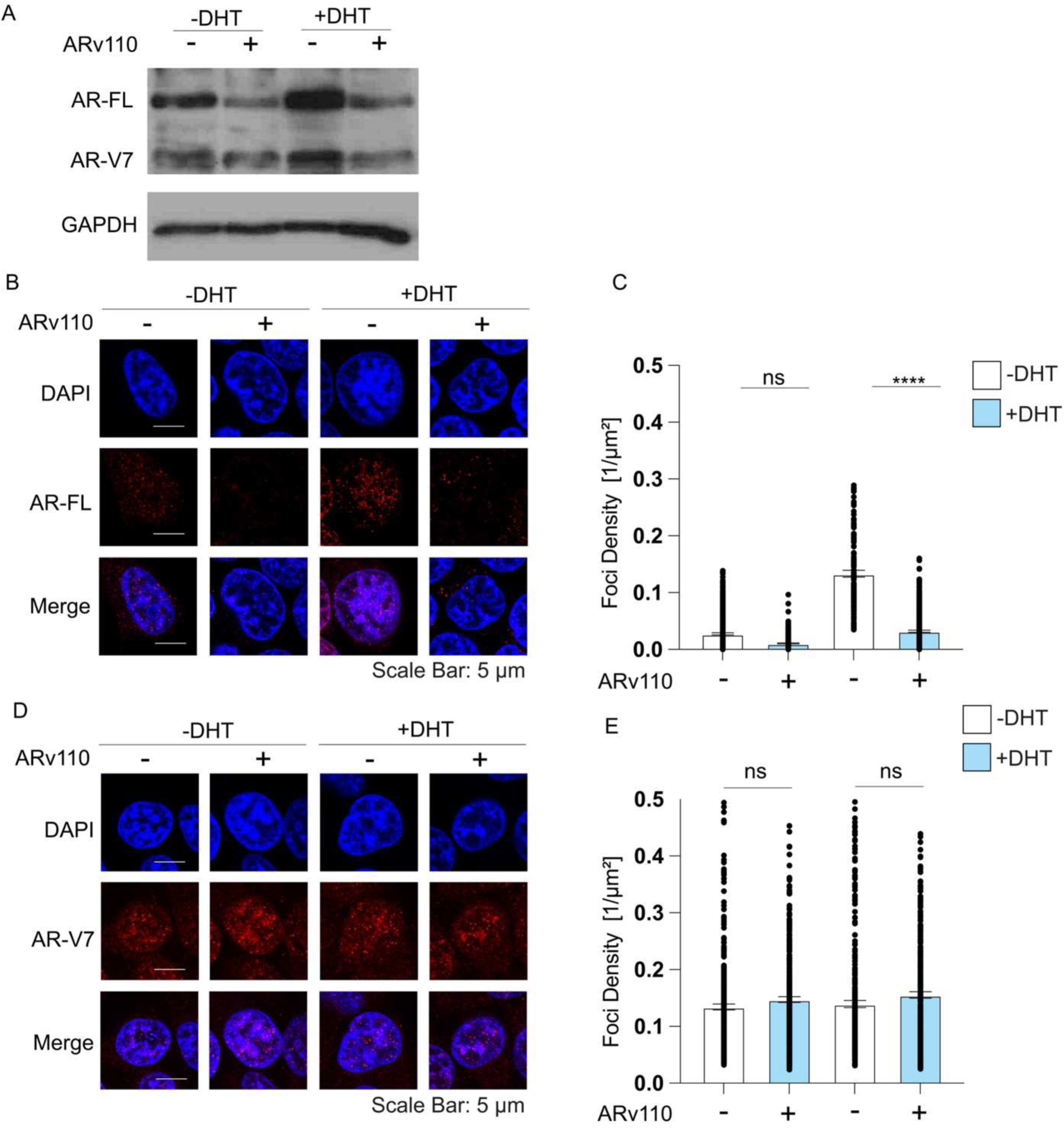
LN95 cells were serum starved for 48 hrs then treated with vehicle or 100 nM ARV110 for 16 hrs. (A) Protein expression levels of endogenous AR-FL and AR-V7 were evaluated by western blot. Endogenous AR-FL (B,C) and AR-V7 (D,E) condensates were detected by immunofluorescence using isoform specific antibodies AR-C19 (sc-815) and AR-V7 (RevMab, 31-1109-00) respectively and analyzed by confocal microscopy (n= three biological replicates and 15 images per replicate). Data are presented as mean ± SEM. Statistical significance was determined using a t-test.

**Supplementary Figure S3:**
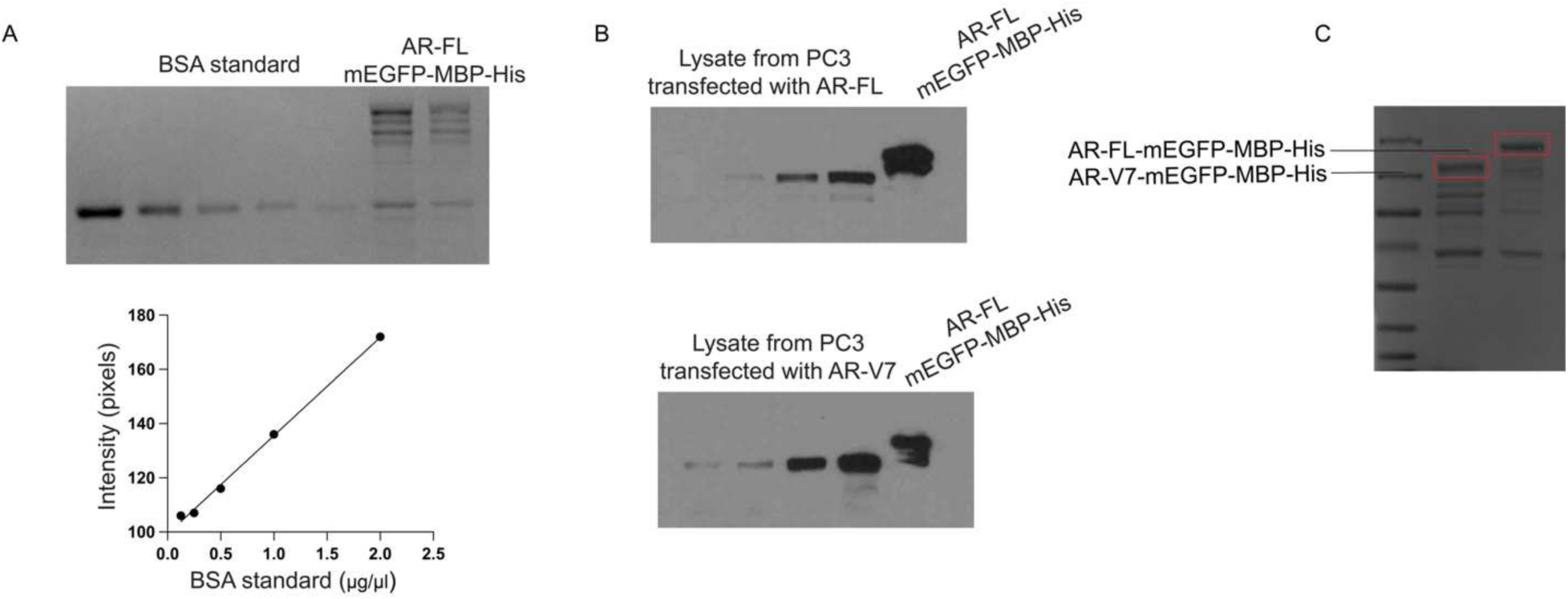
(A) Measuring the concentration of AR-FL-mEGFP-His recombinant protein using a BSA standard titration curve. (B) The known amount of recombinant AR-FL-mEGFP-MBP-His (1μg) was used as a standard to quantify the concentration of AR-FL and AR-V7 in the lysates of PC3 cells. (C) AR-FL-mEGFP-MBP-His and AR-V7-mEGFP-MBP-His recombinant proteins were ran on SDS page gel to ensure the same amount of proteins were used for *in vitro* droplet characterization.

**Supplemental Figure S4.**
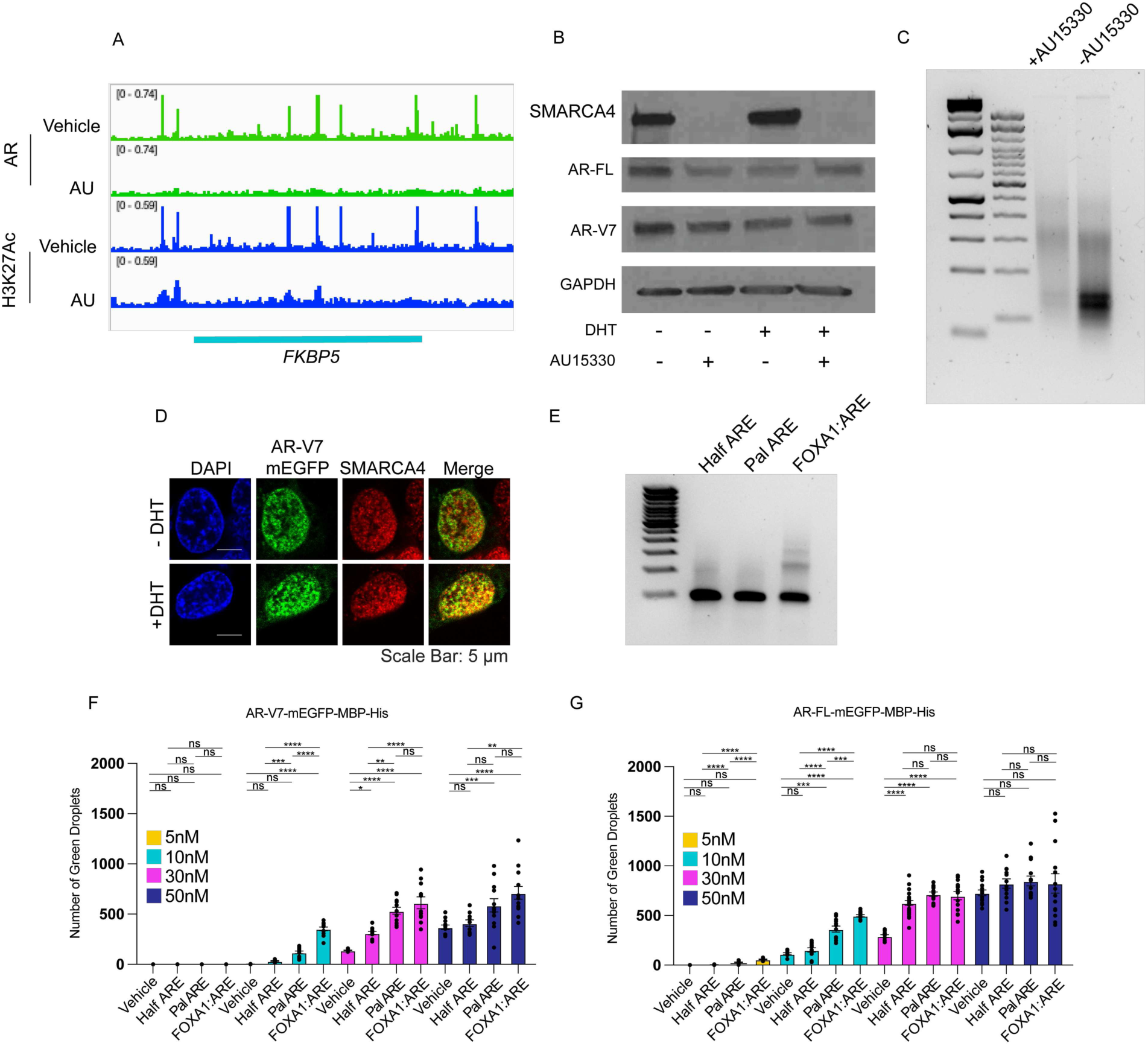
(A) Analysis of ChIP-seq data fin VCaP cells from Xiao, L et al. ^46^ expanding the *FKBP5* gene. (B) Whole cell lysate from treated cells were collected and evaluated by western blot for expression level of target proteins. (C) The chromatin from 22Rv1 cells treated with either AU15330 or vehicle were extracted and digested by MNase and evaluated by agarose gel. (D) 22Rv1 cells were transfected with AR-V7-mEGFP and were cultured in serum starved media. 48 hrs following transfection, cells were examined for colocalization of SMARCA4 with AR-V7-mEGFP by confocal microscopy using antibody specific to SMARCA4. (E) Agarose gel of duplex ROX labeled-half ARE, palindromic ARE and chimeric FOXA1:ARE constructs used in *in vitro assay.* (F-G) Various concentrations of recombinant AR-V7-mEGFP-MBP-His or AR-FL-mEGFP-MBP-His were used *in vitro* to form droplets in the presence of 10% PEG 8000 and either ROX-labeled half, palindromic, or FOXA1:AR chimeric constructs. The number of green droplets was then measured and plotted. The number of green droplets is shown for AR-V7 (F) and AR-FL (G). Data are presented as mean ± SEM. Statistical significance was determined using a one-way ANOVA.

**Supplementary Figure S5.**
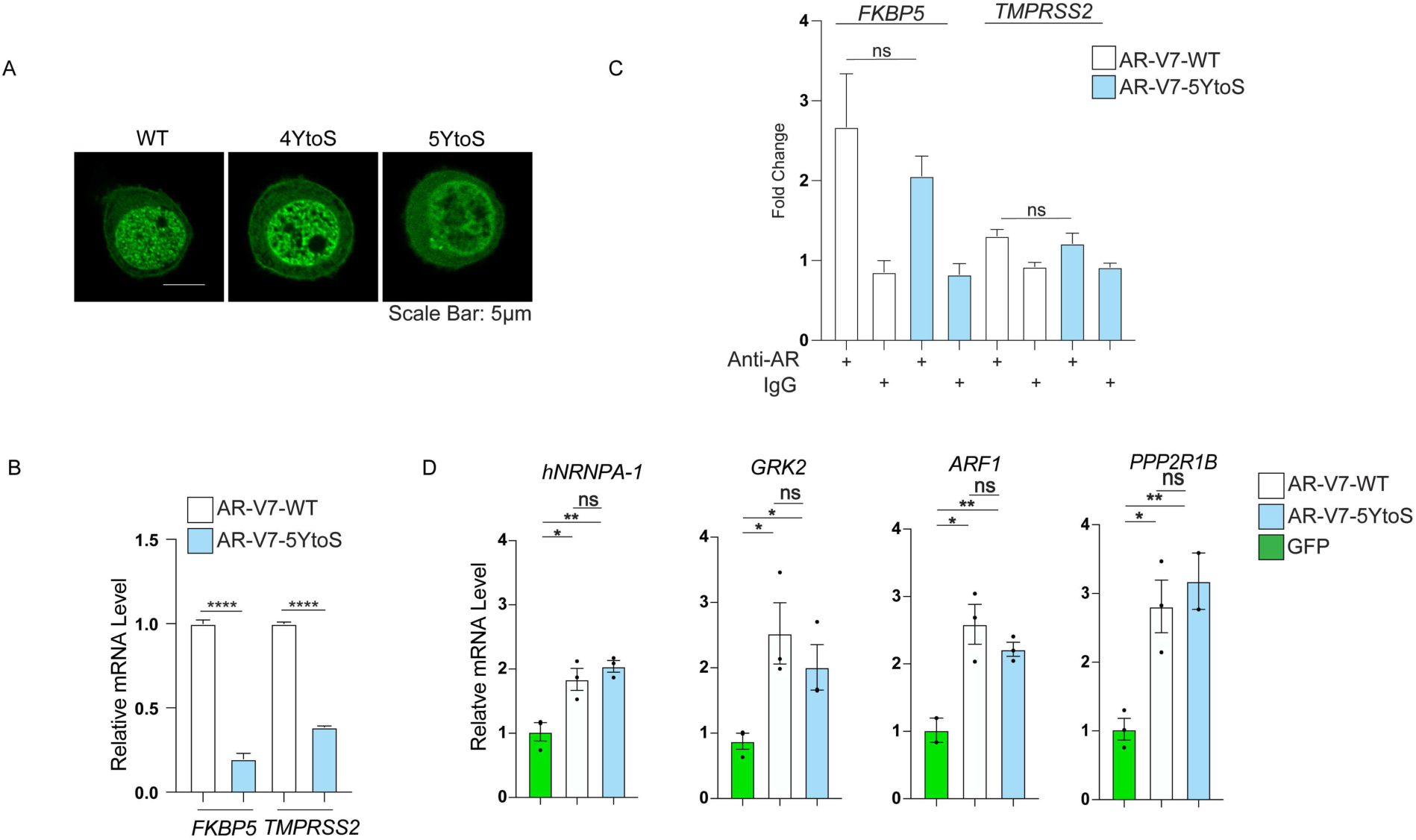
(A) PC3 cells were transfected with wild-type, 4YtoS or 5YtoS AR-V7-mEGFP and were cultured in serum starved media. 48 hrs post-transfection, the cells were then imaged by confocal microscopy. (B) PC3 cells were transfected with AR-V7-WT or AR-V7-5YtoS in serum starved media. 48hrs post transfection, RNA was isolated and synthesized cDNA was used in qPCR. Data are presented as mean ± SEM. Statistical significance was determined using a t-test. (D) PC3 cells were transfected with either AR-V7-mEGFP-WT or AR-V7-mEGFP-5YtoS and serum starved for 48 hrs. The chromatin was next isolated and ChIP assays using anti-AR antibody (AR-441) or IgG (control) were performed. The immunoprecipitated chromatin was then evaluated in qPCR for *FKBP5* and *TMPRSS2* promoter. Data are presented as mean ± SEM. Statistical significance was determined using a t-test. (D) PC3 cells were transfected with either AR-V7-WT, AR-V7-5YtoS or empty-mEGFP plasmids. 48 hrs post-transfection, RNA was isolated, and synthesized cDNA was used in qPCR to assess the expression levels of control genes. Data are presented as mean ± SEM. Statistical significance was determined using a t-test. 22Rv1 cells.

**Supplementary Figure S6.**
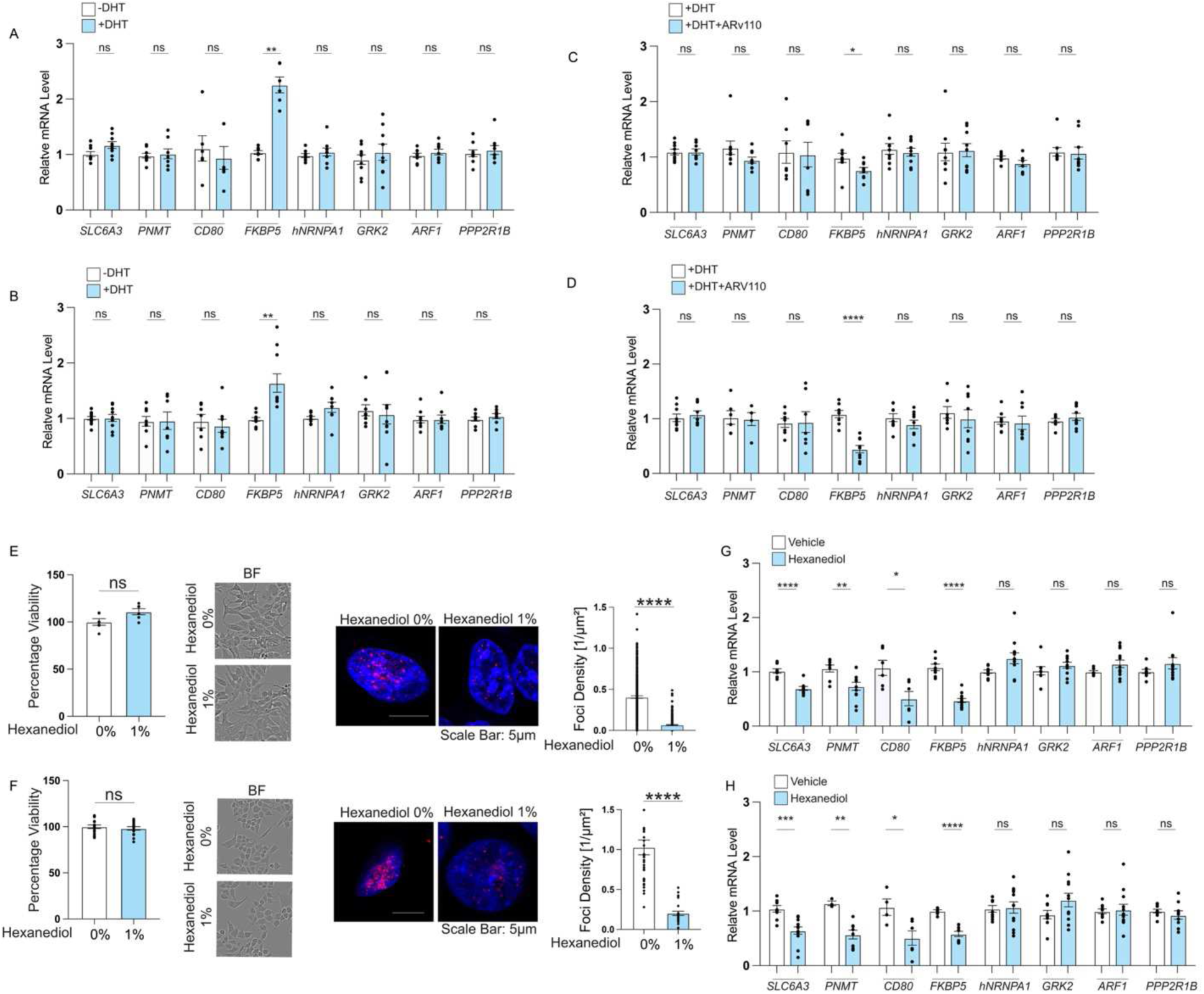
22Rv1 cells (A) and LN95 cells (B) were treated with either vehicle or 1 nM DHT for 2 hrs and the relative expression of target genes were analyzed by qPCR. Data are presented as mean ± SEM. Statistical significance was determined using a t-test. 22Rv1 cells (C) and LN95 cells (D) were treated with either vehicle or 100 nM ARV110 for 16 hrs followed by a stimulation with 1 nM DHT for 2 hrs. The relative expression of target genes was measured using qPCR. Data are presented as mean ± SEM. Statistical significance was determined using a t-test. 22Rv1 cells (E) and LN95 cells (F) were treated with either vehicle or 1% Hexanediol for 30 minutes. The viability was measured using CyQUANT according to manufacturing protocol. The morphology of cells were assed by microscopy (bright field). AR-V7 condensates were detected by immunofluorescence staining using an AR-V7-specific antibody and foci density were measured. 22Rv1 cells (G) and LN95 cells (H) were either vehicle or 1% Hexanediol for 30 minutes. The RNA were then isolated and synthesized cDNA were used in qPCR. Data are presented as mean ± SEM. Statistical significance was determined using a t-test.

## Reference

1. Altmeyer, M. et al. Liquid demixing of intrinsically disordered proteins is seeded by poly(ADP-ribose). Nat Commun 6, 8088 (2015).

2. Banani, S.F., Lee, H.O., Hyman, A.A. & Rosen, M.K. Biomolecular condensates: organizers of cellular biochemistry. Nat Rev Mol Cell Biol 18, 285–298 (2017).

3. Boija, A. et al. Transcription Factors Activate Genes through the Phase-Separation Capacity of Their Activation Domains. Cell 175, 1842–1855 e1816 (2018).

4. Cho, W.K. et al. Mediator and RNA polymerase II clusters associate in transcription-dependent condensates. Science (New York, N.Y.) 361, 412–415 (2018).

5. Kamagata, K. et al. Liquid-like droplet formation by tumor suppressor p53 induced by multivalent electrostatic interactions between two disordered domains. Sci Rep 10, 580 (2020).

6. Tong, X. et al. Liquid-liquid phase separation in tumor biology. Signal Transduct Target Ther 7, 221 (2022).

7. Sharma, R. et al. Liquid condensation of reprogramming factor KLF4 with DNA provides a mechanism for chromatin organization. Nat Commun 12, 5579 (2021).

8. Quail, T. et al. Force generation by protein-DNA co-condensation. Nat Phys 17, 1007-+ (2021).

9. Frank, F., Liu, X. & Ortlund, E.A. Glucocorticoid receptor condensates link DNA-dependent receptor dimerization and transcriptional transactivation. Proc Natl Acad Sci U S A 118 (2021).

10. Yang, J. et al. MYC phase separation selectively modulates the transcriptome. Nat Struct Mol Biol 31, 1567–1579 (2024).

11. Zhang, Y. et al. Nuclear condensates of p300 formed though the structured catalytic core can act as a storage pool of p300 with reduced HAT activity. Nat Commun 12, 4618 (2021).

12. Keenen, M.M. et al. HP1 proteins compact DNA into mechanically and positionally stable phase separated domains. Elife 10 (2021).

13. Bouchard, J.J. et al. Cancer Mutations of the Tumor Suppressor SPOP Disrupt the Formation of Active, Phase-Separated Compartments. Mol Cell 72, 19–36 e18 (2018).

14. Deng, S. et al. RNA m(6)A regulates transcription via DNA demethylation and chromatin accessibility. Nat Genet 54, 1427–1437 (2022).

15. Iannucci, L.F. et al. Cyclic AMP induces reversible EPAC1 condensates that regulate histone transcription. Nat Commun 14, 5521 (2023).

16. Martin, E.W. & Holehouse, A.S. Intrinsically disordered protein regions and phase separation: sequence determinants of assembly or lack thereof. Emerg Top Life Sci 4, 307–329 (2020).

17. Lyons, H. et al. Functional partitioning of transcriptional regulators by patterned charge blocks. Cell 186, 327–345 e328 (2023).

18. Zhang, F. et al. Dynamic phase separation of the androgen receptor and its coactivators key to regulate gene expression. Nucleic Acids Res 51, 99–116 (2023).

19. Ahmed, J., Meszaros, A., Lazar, T. & Tompa, P. DNA-binding domain as the minimal region driving RNA-dependent liquid-liquid phase separation of androgen receptor. Protein Sci 30, 1380–1392 (2021).

20. Basu, S. et al. Rational optimization of a transcription factor activation domain inhibitor. Nat Struct Mol Biol 30, 1958–1969 (2023).

21. Roggero, C.M. et al. A detailed characterization of stepwise activation of the androgen receptor variant 7 in prostate cancer cells. Oncogene 40, 1106–1117 (2021).

22. Xie, J. et al. Targeting androgen receptor phase separation to overcome antiandrogen resistance. Nat Chem Biol 18, 1341–1350 (2022).

23. Viswanathan, S.R. et al. Structural Alterations Driving Castration-Resistant Prostate Cancer Revealed by Linked-Read Genome Sequencing. Cell 174, 433–447 e419 (2018).

24. Zhu, Y. & Luo, J. Regulation of androgen receptor variants in prostate cancer. Asian J Urol 7, 251–257 (2020).

25. Andersen, R.J. et al. Regression of castrate-recurrent prostate cancer by a small-molecule inhibitor of the amino-terminus domain of the androgen receptor. Cancer Cell 17, 535–546 (2010).

26. Lin, T.T. et al. Risk factors for progression to castration-resistant prostate cancer in metastatic prostate cancer patients. J Cancer 10, 5608–5613 (2019).

27. Antonarakis, E.S., Armstrong, A.J., Dehm, S.M. & Luo, J. Androgen receptor variant-driven prostate cancer: clinical implications and therapeutic targeting. Prostate Cancer Prostatic Dis 19, 231–241 (2016).

28. Lallous, N. et al. Functional analysis of androgen receptor mutations that confer anti-androgen resistance identified in circulating cell-free DNA from prostate cancer patients. Genome Biol 17, 10 (2016).

29. Montgomery, R.B. et al. Maintenance of intratumoral androgens in metastatic prostate cancer: a mechanism for castration-resistant tumor growth. Cancer Res 68, 4447–4454 (2008).

30. Watson, P.A., Arora, V.K. & Sawyers, C.L. Emerging mechanisms of resistance to androgen receptor inhibitors in prostate cancer. Nat Rev Cancer 15, 701–711 (2015).

31. Xiang, W. & Wang, S. Therapeutic Strategies to Target the Androgen Receptor. J Med Chem 65, 8772–8797 (2022).

32. Sharp, A. et al. Androgen receptor splice variant-7 expression emerges with castration resistance in prostate cancer. The Journal of clinical investigation 129, 192–208 (2019).

33. Thiyagarajan, T. et al. Inhibiting androgen receptor splice variants with cysteine-selective irreversible covalent inhibitors to treat prostate cancer. Proc Natl Acad Sci U S A 120, e2211832120 (2023).

34. Ehsani, M., David, F.O. & Baniahmad, A. Androgen Receptor-Dependent Mechanisms Mediating Drug Resistance in Prostate Cancer. Cancers (Basel) 13 (2021).

35. Nickols, N.G. et al. MEK-ERK signaling is a therapeutic target in metastatic castration resistant prostate cancer. Prostate Cancer Prostatic Dis 22, 531–538 (2019).

36. Wang, X.S. et al. Characterization of KRAS rearrangements in metastatic prostate cancer. Cancer Discov 1, 35–43 (2011).

37. Weng, C.C. et al. Mutant Kras-induced upregulation of CD24 enhances prostate cancer stemness and bone metastasis. Oncogene 38, 2005–2019 (2019).

38. Hu, R. et al. Distinct transcriptional programs mediated by the ligand-dependent full-length androgen receptor and its splice variants in castration-resistant prostate cancer. Cancer Res 72, 3457–3462 (2012).

39. Ozgun, F. et al. DNA binding alters ARv7 dimer interactions. J Cell Sci 134 (2021).

40. Watson, P.A. et al. Constitutively active androgen receptor splice variants expressed in castration-resistant prostate cancer require full-length androgen receptor. Proc Natl Acad Sci U S A 107, 16759–16765 (2010).

41. Xu, D. et al. Androgen Receptor Splice Variants Dimerize to Transactivate Target Genes. Cancer Res 75, 3663–3671 (2015).

42. Cai, L. et al. ZFX Mediates Non-canonical Oncogenic Functions of the Androgen Receptor Splice Variant 7 in Castrate-Resistant Prostate Cancer. Mol Cell 72, 341–354 e346 (2018).

43. Chen, Z. et al. Diverse AR-V7 cistromes in castration-resistant prostate cancer are governed by HoxB13. Proc Natl Acad Sci U S A 115, 6810–6815 (2018).

44. Liang, J. et al. Androgen receptor splice variant 7 functions independently of the full length receptor in prostate cancer cells. Cancer Lett 519, 172–184 (2021).

45. Gao, X.e.a. Phase 1/2 study of ARV-110, an androgen receptor (AR) PROTAC degrader, in metastatic castration-resistant prostate cancer (mCRPC). Clinical Oncology 40 (2022).

46. Xiao, L. et al. Targeting SWI/SNF ATPases in enhancer-addicted prostate cancer. Nature 601, 434–439 (2022).

47. Parolia, A. et al. NSD2 is a requisite subunit of the AR/FOXA1 neo-enhanceosome in promoting prostate tumorigenesis. Nat Genet (2024).

48. Chen, C., Fu, G., Guo, Q., Xue, S. & Luo, S.Z. Phase separation of p53 induced by its unstructured basic region and prevented by oncogenic mutations in tetramerization domain. Int J Biol Macromol 222, 207–216 (2022).

49. Sugiura, M. et al. Identification of AR-V7 downstream genes commonly targeted by AR/AR-V7 and specifically targeted by AR-V7 in castration resistant prostate cancer. Transl Oncol 14, 100915 (2021).

50. Davis, J.E. et al. ARF1 promotes prostate tumorigenesis via targeting oncogenic MAPK signaling. Oncotarget 7, 39834–39845 (2016).

51. Pandey, P. et al. Impaired expression of protein phosphatase 2A subunits enhances metastatic potential of human prostate cancer cells through activation of AKT pathway. Br J Cancer 108, 2590–2600 (2013).

52. Zhang, M. et al. Targeting the Lnc-OPHN1-5/androgen receptor/hnRNPA1 complex increases Enzalutamide sensitivity to better suppress prostate cancer progression. Cell Death Dis 12, 855 (2021).

53. Basil, P. et al. Cistrome and transcriptome analysis identifies unique androgen receptor (AR) and AR-V7 splice variant chromatin binding and transcriptional activities. Sci Rep 12, 5351 (2022).

54. Krause, W.C., Shafi, A.A., Nakka, M. & Weigel, N.L. Androgen receptor and its splice variant, AR-V7, differentially regulate FOXA1 sensitive genes in LNCaP prostate cancer cells. Int J Biochem Cell Biol 54, 49–59 (2014).

55. Luo, J. et al. Role of Androgen Receptor Variants in Prostate Cancer: Report from the 2017 Mission Androgen Receptor Variants Meeting. Eur Urol 73, 715–723 (2018).

56. Monaghan, A.E. & McEwan, I.J. A sting in the tail: the N-terminal domain of the androgen receptor as a drug target. Asian J Androl 18, 687–694 (2016).

57. Nagandla, H. et al. Isoform-specific Activities of Androgen Receptor and its Splice Variants in Prostate Cancer Cells. Endocrinology 162 (2021).

58. Banuelos, C.A. et al. Sintokamide A Is a Novel Antagonist of Androgen Receptor That Uniquely Binds Activation Function-1 in Its Amino-terminal Domain. J Biol Chem 291, 22231–22243 (2016).

59. Shimizu, Y. et al. Androgen Receptor Splice Variant 7 Drives the Growth of Castration Resistant Prostate Cancer without Being Involved in the Efficacy of Taxane Chemotherapy. J Clin Med 7 (2018).

60. Zhang, X. et al. Androgen receptor variants occur frequently in castration resistant prostate cancer metastases. PLoS One 6, e27970 (2011).

61. Shrinivas, K. et al. Enhancer Features that Drive Formation of Transcriptional Condensates. Mol Cell 75, 549–561 e547 (2019).

62. Chen, L. et al. Hormone-induced enhancer assembly requires an optimal level of hormone receptor multivalent interactions. Mol Cell 83, 3438–3456 e3412 (2023).

63. Deshpande, P., Prentice, E., Vidal Ceballos, A., Casaccia, P. & Elbaum-Garfinkle, S. Epigenetic marks uniquely tune the material properties of HP1alpha condensates. Biophys J 123, 1508–1518 (2024).

64. Cato, L. et al. ARv7 Represses Tumor-Suppressor Genes in Castration-Resistant Prostate Cancer. Cancer Cell 35, 401–413 e406 (2019).

65. Han, D. et al. Androgen receptor splice variants drive castration-resistant prostate cancer metastasis by activating distinct transcriptional programs. J Clin Invest 134 (2024).

66. Du, M. et al. Direct observation of a condensate effect on super-enhancer controlled gene bursting. Cell 187, 2595–2598 (2024).

67. Miller, K.J. et al. A compendium of Androgen Receptor Variant 7 target genes and their role in Castration Resistant Prostate Cancer. Front Oncol 13, 1129140 (2023).

68. Liu, J. et al. Targeting NSD2-mediated SRC-3 liquid-liquid phase separation sensitizes bortezomib treatment in multiple myeloma. Nat Commun 12, 1022 (2021).

69. Wang, S. et al. ALDH1A3 serves as a predictor for castration resistance in prostate cancer patients. BMC Cancer 20, 387 (2020).

70. Liao, C. et al. SPINKs in Tumors: Potential Therapeutic Targets. Front Oncol 12, 833741 (2022).

71. Jansen, F.H. et al. Profiling of antibody production against xenograft-released proteins by protein microarrays discovers prostate cancer markers. J Proteome Res 11, 728–735 (2012).

72. Taichman, R.S. et al. GAS6 receptor status is associated with dormancy and bone metastatic tumor formation. PLoS One 8, e61873 (2013).

73. Yang, Q. et al. MAZ promotes prostate cancer bone metastasis through transcriptionally activating the KRas-dependent RalGEFs pathway. J Exp Clin Cancer Res 38, 391 (2019).

74. Pomp, W., Meeussen, J.V.W. & Lenstra, T.L. Transcription factor exchange enables prolonged transcriptional bursts. Mol Cell 84, 1036–1048 e1039 (2024).

75. Trojanowski, J. et al. Transcription activation is enhanced by multivalent interactions independent of phase separation. Mol Cell 82, 1878–1893 e1810 (2022).

